# Charting spatial ligand-target activity using Renoir

**DOI:** 10.1101/2023.04.14.536833

**Authors:** Narein Rao, Tanush Kumar, Rhea Pai, Archita Mishra, Florent Ginhoux, Jerry Chan, Ankur Sharma, Hamim Zafar

**Author notes:** Corresponding Author* (A.S.), (H.Z.).

## Abstract

The advancement of single-cell RNA sequencing (scRNA-seq) and spatial transcriptomics has enabled the inference of cellular interactions in a tissue microenvironment. Despite the development of cell-cell interaction inference methods, there is a lack of methods capable of mapping the influence of ligands on downstream target genes across a spatial topology with specific cell type composition, with the potential to shed light on niche-specific relationship between ligands and their downstream targets. Here we present Renoir for charting the ligand-target activities across a spatial topology and delineating spatial communication niches harboring specific ligand-target activities. Renoir also spatially maps pathway-level activity of ligand-target genesets and identifies domain-specific ligand-target activities. Across spatial datasets with varying resolution (spot to single-cell) ranging from development to disease, Renoir inferred cellular niches with distinct ligand-target interactions, spatially mapped pathway activities, and identified context-specific novel cell-cell interactions. Renoir uncovers biological insights and therapeutically-relevant cellular crosstalk from spatial transcriptomics data.

## Introduction

Coordination among the cells drive various functions in a multicellular tissue where cellular decision making is influenced by interactions with the environment composed of extrinsic stimuli and other cells. These interactions also known as cell–cell communication (CCC) play a crucial role in tissue development, biological processes and functions (1). Recently emerged single-cell RNA sequencing (scRNA-seq) technologies have made it possible to analyze tissue architecture at the resolution of individual cells (2; 3) and have shown great potential in investigating CCC (4). Computational methods have been developed for inferring cell-cell communication and mediating ligand-receptor (LR) pairs from scRNA-seq datasets by utilising ligand-receptor databases (5; 6). These methods including CellChat (6), CellphoneDB (5), iCELLNET (7), NATMI (8), scTensor (9), and SingleCellSignalR (10) mostly rely on defining functions based on the expression of ligand and receptor pair. Beyond ligand-receptor interactions, methods such as NicheNet (11), Cytotalk (12), SoptSC (13) and CCCExplorer (14) focused on inferring the effects of ligand–receptor interactions on possible downstream target gene expression in the receiving cells. While these methods have provided biological insights based on single-cell datasets, they often generate false-positives by failing to account for the spatial context of cell type arrangement in the tissue, as the spatial location information is not captured in scRNA-seq data. Retaining the spatial context is of paramount importance in CCC inference as cellular communications are often spatially constrained where signalling pathways are activated when ligands diffuse from sender cells to neighbouring receiver cells in close proximity (9).

Recent breakthroughs in sequencing (15; 16) and imaging-based (17) spatial transcriptomic technologies have enabled the measurement of gene expression profiles at varying cellular resolutions from single-cells to spots consisting of multiple or fractions of cells while preserving the spatial context information for the cells or spots. Such datasets have been utilized for spatially mapping cells from scRNA-seq data for inferring ligand-target interactions (e.g., SpaOTsc (18)). Moreover, CCC inference methods that specifically use spatial data have also been developed. Methods such as Giotto (19) and NCEM (20) use a spatial neighbor graph for inferring cell-cell interactions; CellPhoneDB v3 (21) uses the spatial information to define microenvironments within which interactions are restricted; stLearn (22) and SpatialDM (23) rely on the co-expression of ligand and receptor genes in relation to the spatial location diversity; SVCA (24), MISTy (25) and COMMOT (26) use probabilistic model, machine learning model and optimal transport respectively for identifying spatially restricted ligand-receptor interactions. However, most of these methods focus on ligand-receptor interactions, they do not examine the effect of the cell-cell interaction on the down-stream target gene expression. Moreover, most of these methods aim to uncover the interacting cell types globally and thus may be less sensitive in identifying local interactions. While a handful of methods including COMMOT, stLearn and SpatialDM can compute ligand-receptor interactions at the resolution of a spatial location, they fail to quantify the activity of ligand-target gene pairs at the same resolution given a specific spatial context with a fine-grained spatial distribution of cell types. Furthermore, the existing methods employing low-resolution spatial datasets (e.g., 10X Visium) cannot quantify cell type-specific interactions from the mixture of cell types present in a spot. Given that the existing computational methods were developed based on low-resolution spatial datasets, they may provide sub-optimal results for the recent single-cell resolution spatial technologies including NanoString CosMx and 10X Xenium. There is a critical requirement for reliable computational approaches that can elucidate the probable influence of the ligands on target gene expressions in the receiving cell in a particular spatial context with unique cell type composition.

To address these challenges, we introduce Renoir (ligand-ta<u>R</u>g<u>e</u>t i<u>n</u>teractions acr<u>o</u>ss spat<u>i</u>al topog<u>r</u>aphy), an end-to-end computational framework for spatially charting the ligand-target activities and inferring spatial communication niches that harbor specific ligand-target activities and cell type composition. Renoir requires either single-cell resolution spatial transcriptomic data or matched low-resolution spatial transcriptomic data and scRNA-seq data from the same tissue in order to delineate the spatial influence between possible ligand-target pairs curated from an existing database (11). Therefore, Renoir allows one to determine which ligands lead to the activation of which target genes in a given spatial context. For each curated ligand-target pair, Renoir quantifies a neighborhood activity score for each spatial location of the spatial transcriptomics data by taking into account the cell type abundances, cell type-specific mRNA abundance values of the ligand and target genes, expression of suitable receptors for the ligand in the cell type harboring target gene, cell type-specific gene entropy measure and a cell type-specific mutual information between the ligand-target pairing as obtained from single-cell resolution spatial data or scRNAseq data. The neighborhood activity score computed by Renoir denotes the strength of the ligand-target activity within the context of a spot’s neighbourhood. In addition to mapping potential ligand-target relationships throughout a spatial topology, the neighborhood scores computed by Renoir further enables a wide variety of downstream analyses. Using the neighbourhood scores, Renoir can infer communication niches, which constitute tissue microenvironments harboring specific ligand-target activities pertaining to major co-localized cell types. The neighborhood scores also allow for the spatial mapping of the total activity of gene sets corresponding to well-established signaling pathways as well as inferred denovo through unsupervised clustering. Renoir can also identify the major ligands active in a communication niche and rank them based on their cumulative activity across the target genes expressed in the niche.

Using a wide range of synthetic datasets generated across a variety of tissue types (brain, cancer and intestine) as well as real datasets, we demonstrated that Renoir outperforms the state-of-the-art methods in inferring spatial activity of ligand-target pairs and communication niches. To demonstrate the utility of Renoir in uncovering spatial map of ligand-target activities across tissue types, we applied Renoir to three spatial transcriptomic datasets ranging from development to disease, including adult mouse brain, human triple negative breast cancer (TNBC), and human fetal liver, for which Renoir uncovered distinct spatial domains characterized by specific ligand-target activities. Renoir characterized the interactions of regional astrocyte subtypes with other spatially co-localized cell types in the mouse brain, identified several spatially restricted interactions between cancer, stromal and immune cell populations in TNBC, and uncovered hepatocyte-macrophage interactions in developing fetal liver. To further demonstrate Renoir’s ability to handle single-cell resolution spatial datasets, we applied it on a NanoString CosMx dataset from hepatocellular carcinoma for which Renoir identified known and novel onco-fetal interactions.

## Results

### Charting spatial ligand-target interactions using Renoir

In order to build a comprehensive computation framework to investigate spatial cell-cell interaction we developed Renoir. As shown in Figure 1, Renoir is an end-to-end computational framework for exploring the spatial map of ligand-target activities either by integrating spatial transcriptomics and scRNA-seq datasets from the same tissue or by employing single-cell resolution spatial transcriptomics data. Renoir first curates a set of ligand-target pairs using NATMI’s ConnectomeDB (8) and NicheNet (11). For a low-resolution spatial transcriptomic dataset (e.g., 10X Visium), using a well-annotated scRNA-seq data, Renoir quantifies a neighborhood activity score for each curated ligand-target pair for each spot in the spatial transcriptomic data. For a single-cell resolution spatial transcriptomic data, Renoir quantifies the neighborhood activity score for each curated ligand-target pair for each cell with a spatial context. The quantification of the neighborhood activity score for a specific ligand-target pair for a particular spot is performed using the proportions and cell type-specific mRNA abundances present in each spot/cell within the defined neighborhood of the spot/cell (see Methods for details). The activity between a ligand-target pair for the given spot and another spot in the neighborhood is scored based on the cell type–specific mRNA abundances of the two genes weighted by the cell type proportions as well as the mutual information between the two genes across the cell types present within the two spots. Thus, Renoir generates a novel neighborhood activity score matrix (of dimensions number of ligand-target pairs × number of spots). Using the ligand-target neighborhood activity score matrix, Renoir performs several downstream analyses. First of all, Renoir infers spatial communication domains through unsupervised clustering of spots based on ligand-target neighborhood activity scores and determines communication domain-specific ligand-target activities. Moreover, Renoir can infer cell type-specific ligand-target activities within a communication domain. Renoir further provides an algorithm for ranking the ligands within a communication domain based on their cumulative activity across target genes expressed by the major cell types in the domain. Renoir also provides a mapping of pathway-level activity across spots and communication domains. Finally, Renoir encompasses a visualization module for the visualisation of ligand-target activity across spots as well as a Sankey plot that provides an overview of user-specified ligand-target activities across cell types and communication domains.

**Figure 1.**
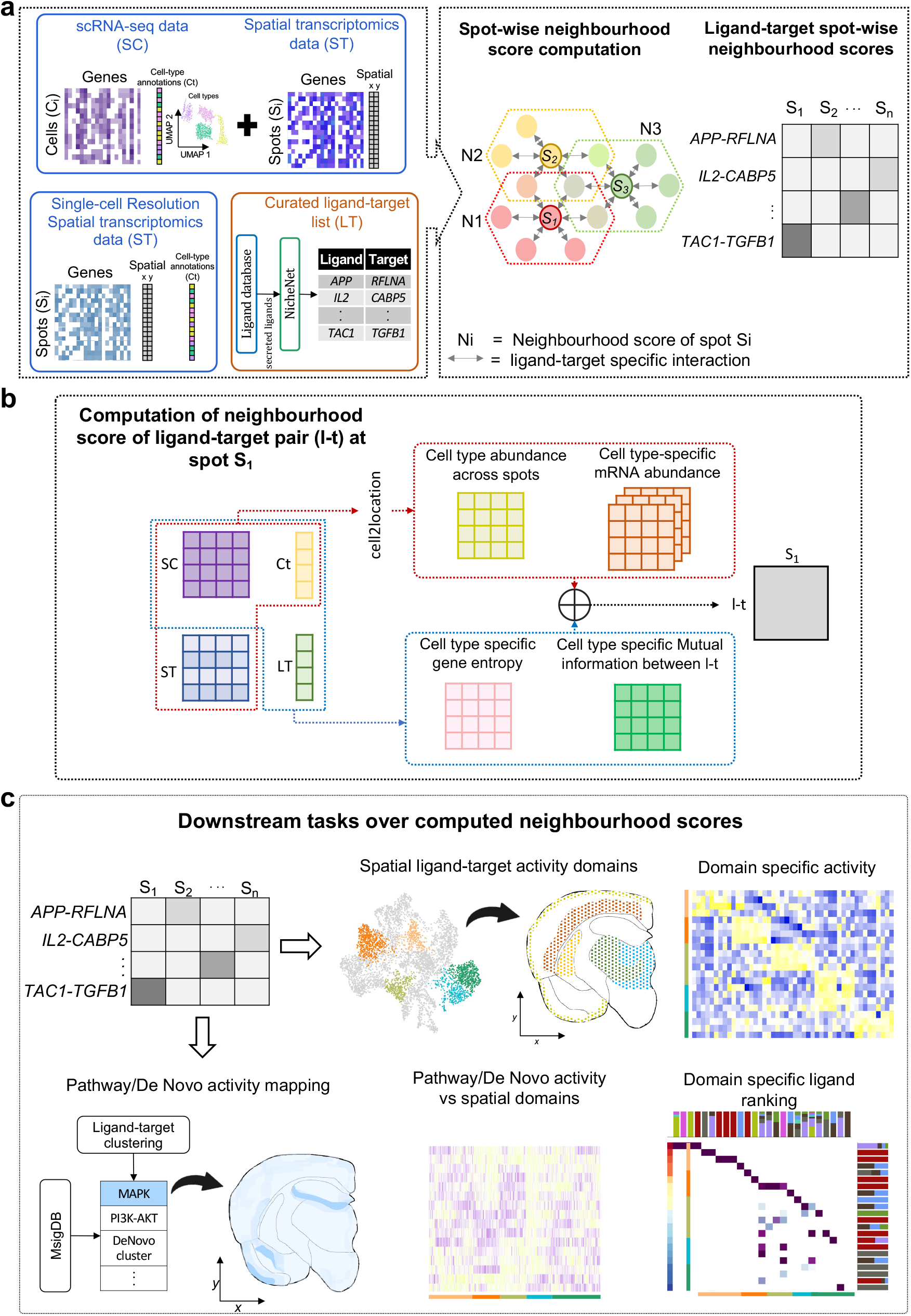
Overview of Renoir. (a) Renoir utilizes either spot-resolution spatial transcriptomic data and cell type-annotated scRNA-seq data from the same tissue or single-cell resolution spatial transcriptomic data to infer spatial neighborhood activity scores for a set of curated ligand-target pairs at each spot. (b) To estimate the neighborhood activity score for a ligand-target pair *l* − *t* at a given spot *S*1; Renoir utilizes cell type abundance, cell type-specific mRNA abundance for each spot as inferred by a cell type deconvolution module, expression of the receptor at the target cell types, gene entropy and mutual information between l-t specific to cell types. (c) Renoir performs various downstream tasks based on the ligand-target neighborhood scores - inference of spatial communication domains and domain-specific ligand-target activities, spatial mapping of pathway activity and ranking of ligand activity.

### Renoir outperforms state-of-the-art methods in inferring ligand-target interactions

We first evaluated Renoir’s performance in inferring the spatial activity of ligand-target pairs using simulated datasets generated using a semi-synthetic enrichment-based simulation workflow (Fig. 2a, see Methods for details). Specifically, we started with paired single-cell and spatial datasets based on which we simulated new spatial datasets where activity of a ligand-target pair was simulated in reference spatial locations (positive reference spots) through the enrichment of cell types associated with the ligand and target gene. Moreover, the simulated datasets also contained spatial locations which did not harbor any activity between the ligand and target (negative reference spots). To comprehensively cover the range of interactions across tissue types, we used three reference matched scRNA-seq and ST datasets from human intestine (27), triple negative breast cancer (TNBC) (28) and dorsolateral prefrontal cortex (DLPFC) (29) for the simulation. The simulation workflow as described in Figure 2a was applied to generate 43, 15 and 26 simulated datasets corresponding to the human intestine, TNBC and DLPFC datasets respectively, where, each simulated dataset enriched a specific ligand-target interaction. Since Renoir quantifies ligand-target activity at a specific spatial context and is the only method (to the best of our knowledge) to do so contrary to the other cell-cell interaction methods, which infer ligand-receptor interactions by employing the spatial data, as competitor methods, we selected state-of-the-art ligand-receptor interaction inference methods COMMOT (26), stLearn (30) and SpatialDM (23). These three methods are capable of quantifying ligand-receptor interactions at the resolution of spatial spots and we adopted them for quantifying ligand-target activity from our simulated datasets. The performance of each method was evaluated based on a spatial activity accuracy metric that computes the Pearson correlation of the spot-level ligand-target activity score measured by a method with the true simulated activity of the ligand-target pair in the positive and negative reference spatial locations (see Method for details). In addition, we also compared the methods in terms of precision and recall (see Methods) for the ligand-target pairs to evaluate their ability to distinguish the spots harboring ligand-target interactions from the spots devoid of such interactions.

**Figure 2.**
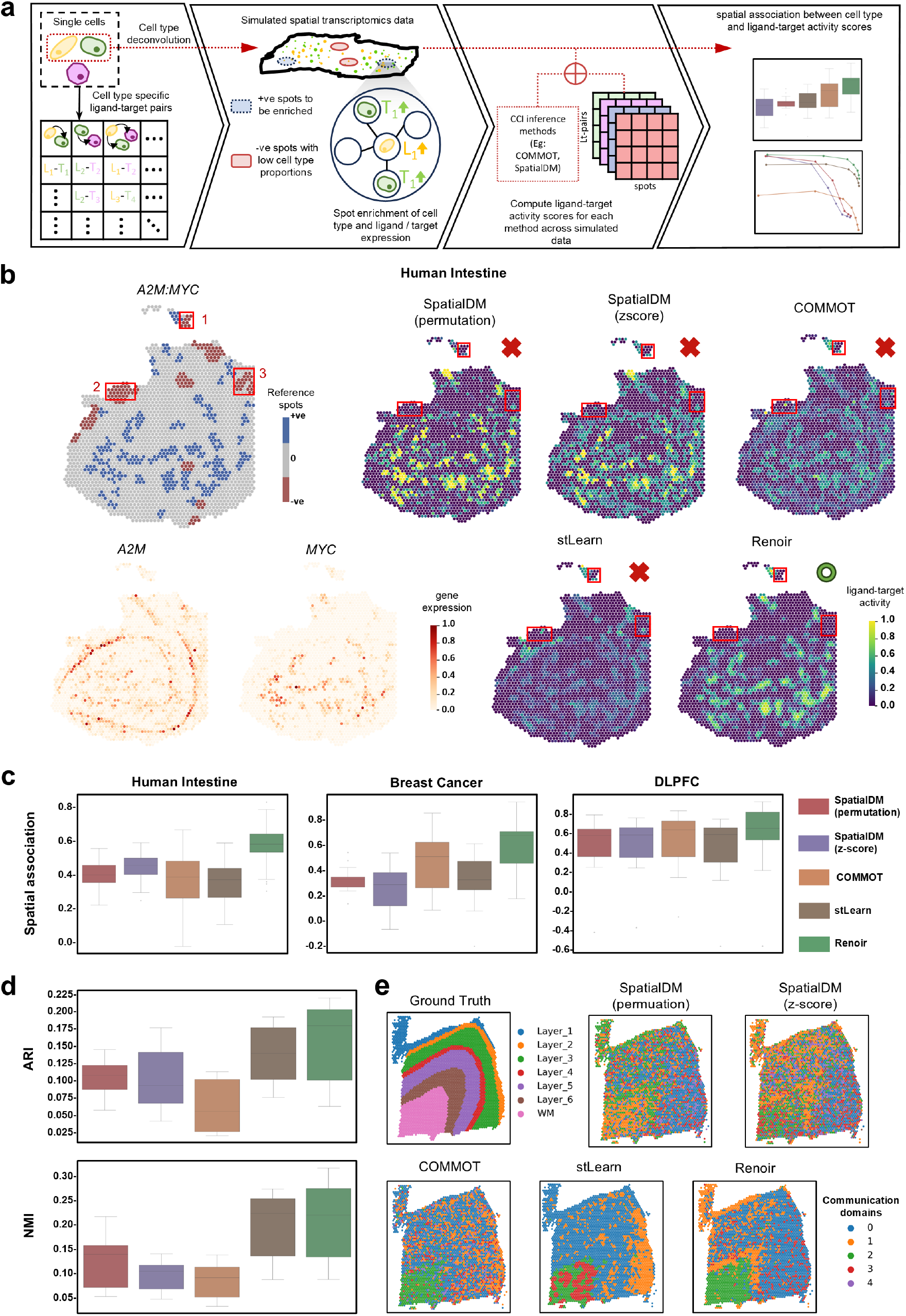
Renoir outperforms state-of-the-art CCC inference methods in inferring ligand-target activities. (a) Overview of the simulation strategy used for benchmarking. (b) Comparison of Renoir, SpatialDM, COMMOT, and stLearn for the simulated dataset for the ligand-target pair *A2M-MYC* simulated based on the human intestine dataset. Positive and negative reference spots are shown in the tissue. Regions 1, 2 and 3 contain negative spots for which SpatialDM, COMMOT, and stLearn falesly inferred activity of the pair but Renoir correctly inferred the absence of activity. (c) Comparison of Renoir, SpatialDM, COMMOT, and stLearn in terms of spatial activity accuracy over all the ligand-target pairs across the tissue types. (d) ARI and NMI comparison of the spatial communication domains computed based on the ligand-target activity scores inferred by Renoir, SpatialDM, COMMOT, and stLearn. (e) Qualitative comparison of the spatial communication domains inferred by Renoir, SpatialDM, COMMOT, and stLearn for DLPFC sample 151673.

In each simulated dataset across the tissue types, Renoir was able to capture the ligand-target interactions more precisely. To illustrate, we consider the ligand-target pair *A2M-MYC*, simulated from the human intestine dataset (Fig. 2b), where the positive (blue) and negative (red) reference spots included enrichment and abatement of said ligand-target pair interaction respectively as is also evident from the spatial expression plot of the two genes. Amongst the positive spots, we observed Renoir’s ligand-target activity scores to truly reflect the enriched spots with high scores as compared to COMMOT and stLearn. While SpatialDM had high scores for positive spots, it also had scores all across the tissue and failed to differentiate the positive and negative spots appropriately. Renoir was also able to correctly quantify the absence of ligand-target interactions in the negative reference spots. In comparison, all three competitor methods showed false-positive activity of the gene pair in multiple negative spots (red bounded regions labeled 1, 2 and 3) with near obsolete *A2M* and *MYC* expression. Renoir is capable of computing the activity of a ligand-target by considering the expression of a suitable receptor at the receiving cell. To showcase this, we considered the pair *IFNG-JCHAIN*, simulated using the TNBC dataset, where *IFNG* was associated with Natural Killer T Cells (NKT Cells) and CD8^+^ T Cells; and *JCHAIN* was associated with B Cells and Dendritic Cells (DCs). However, B cells do not express a receptor for *IFNG* (Supplementary Fig. 1a) and thus the *IFNG* secreted from NKT Cells or CD8^+^ T Cells cannot activate *JCHAIN* in B cells. Therefore, the positive reference spots harbored interactions with DCs as the target cell type whereas the negative spots contained the enrichment of B cells as the target cell type (see Methods section for more details). In this scenario, by accounting for the presence of suitable receptors while calculating ligand-target activity scores, Renoir correctly scored the negative spots with negligible scores; whereas other methods failed to consider such scenarios and therefore incorrectly gave high scores to the negative spots as depicted by the red bounded regions (Supplementary Fig. 1a). Similarly, for the DLPFC dataset (e.g., for the pair *NPY:CARTPT*) also, Renoir’s spatial scores for the ligand-target activity were more precise as compared to the other methods (Supplementary Fig. 1b). SpatialDM and COMMOT had less distinction between the positive and non-positive spots and stLearn failed to capture the activity pattern due to the high range of *NPY* expression and relatively low range of *CARTPT* expression. Quantitatively, Renoir outperformed other CCC inference methods in inferring spatial ligand-target activity by achieving 45.8, 98.5, and 12.4% improvement in the median spatial activity accuracy across the human intestine, TNBC and DLPFC datasets (Fig. 2c) indicating Renoir’s superiority in capturing the spatial variation of the ligand-target activity. Furthermore, in terms of average precision and recall over the ligand-target pairs, Renoir outperformed SpatialDM, COMMOT, and stLearn across the tissue types across different threshold values used for calculating precision and recall (Supplementary Fig. 1c). For varying threshold values, Renoir was able to consistently maintain high precision and recall values across the tissue types. In comparison, the precision and recall values for the other methods varied across datasets, with a method having high precision-recall for one dataset but low precision-recall for another.

Finally, we evaluated the accuracy of the spatial communications domains inferred based on the ligand-target interaction scores inferred by a CCC inference method. For this, we utilized 10X Visium datasets from 12 dorsolateral prefrontal cortex (DLPFC) samples (29), in which the authors manually annotated six cortical layers and white matter for each sample based on cytoarchitecture and selected marker genes. Considering the manual annotations as ground truth, we evaluated the utility of the ligand-target scores inferred by Renoir in identifying the layers and compared its performance against that of COMMOT, SpatialDM and stLearn. Each method was evaluated based on adjusted Rand index (ARI) and normalized mutual information (NMI) scores calculated by comparing the communication domains inferred by the method against the ground truth annotations over all 12 DLPFC samples (see Methods for details). Renoir achieved the highest ARI and NMI scores considering all the samples and outperformed all the other methods (Fig. 2d) indicating that it is able to capture spatial variation and architecture better than the other CCC inference methods. Figure 2e shows how the communication domains inferred by Renoir captured the underlying spatial structure for the sample 151675 better than the other methods for which the inferred communication domains were much noisier.

### Renoir identifies distinct communication domains in Mouse brain

To demonstrate Renoir’s ability to uncover region-specific ligand-target interactions, we first applied Renoir to 10X Visium data from two Mouse brain sections (samples S1 and S2) and its corresponding single nuclei data (31) (see data availability). Renoir identified 9409 (S1) and 9405 (S2) ligand-target pairs that were expressed in this dataset and quantified their spatial neighborhood activity scores for both samples. Based on the ligand-target neighborhood activity scores, Renoir clustered the ST spots across the two tissue sections and identified 19 spatial communication domains which had distinct latent space embeddings (Fig. 3a) and were similarly distributed across the sections (Fig. 3b). We further quantified the distribution of the spatial communication domains across different anatomical brain regions and found that the domains across both samples primarily corresponded to the same anatomical regions and were highly concordant across the two samples in terms of their distribution in anatomical brain regions (Fig. 3c).

**Figure 3.**
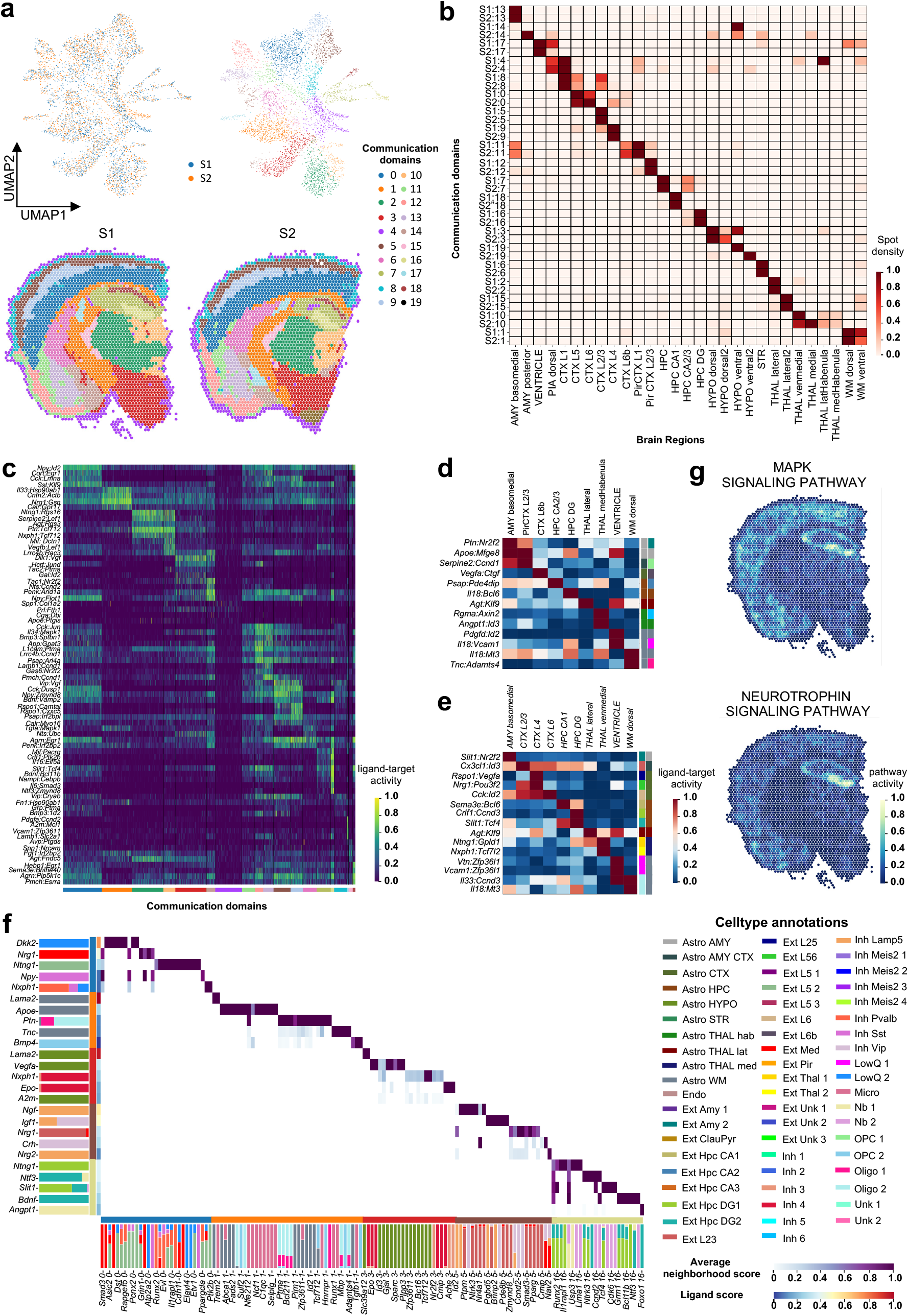
Renoir identifies distinct communication domains in mouse brain. (a) Spatial communication domains with distinct latent embeddings inferred by Renoir based on ligand-target neighborhood scores for two mouse brain sections. (b) Distribution of spots in spatial communication domains across different anatomical regions of the mouse brain. The rows correspond to spatial communication domains (in both S1 and S2) and columns correspond to known brain regions. Each cell represents the proportion of spots of a communication domain in the corresponding anatomical brain region. (c) Differentially active ligand-target interactions across the communication domains for sample S1. (d) - (e) Characterization of ligand-target interactions between regional astrocyte subtypes and other co-localized cell types in sample S1. (d) Ligands expressed by regional astrocyte subtypes activating target genes in other cell types. (e) Target genes in regional astrocyte subtypes activated by ligands expressed by other co-localized cell types. (f) Communication domain-specific ranking of ligands based on cumulative activities over target genes expressed by major cell types in the domain for sample S1. Top three ligands for each of eight communication domains are represented. Stacked color bars represent the cell types that express the ligand (right) and target (top).(g) Spatial map of MAPK SIGNALING and NEUROTROPHIN SIGNALING pathway activity across the spots in sample S1.

The communication domains were found to harbor diverse cell types that were spatially localized and abundance pattern concordant across the replicates (Supplementary Fig. 2a-b). The domains further harbored differential activity of specific ligand-target pairs (Fig. 2d, Supplementary Fig. 3b-c). For instance, domain 1 specific to white matter exhibited *Il33* and *Csf1* activity which are known to be required for microglia development and maintenance in white matter (32; 33) and domain 3 specific to the hypothalamus exhibited *Dlk1* activity known to be expressed in hypothalamic arginine vasopressin and oxytocin neurons (34). We also noticed the existence of ligand or target-specific activity pertaining to distinct anatomical regions, wherein the ligand or target was known to be abundant within the brain regions corresponding to the domain. For example, domain 9 exhibidt high activity of *Rspo1*, which is known to be highly expressed within the cortex (35; 36), domain 7 exhibited high activity of hippocampal marker *Crlf1*, and domain 13 exhibited high activity of *Ccnd1* and *Nr2f2*, which are known to be highly expressed within the Amygdala region (37; 38). Supplementary Table 1 lists the domain-specific ligand-target interactions with literature support for the associated brain region.

Kleshchevnikov et al. (31) identified and molecularly characterized 10 regional astrocyte subtypes using this spatial and matched single-cell data. However, interactions of these regional astrocyte subtypes with other spatially co-localized cell types were not explored. We applied Renoir to characterize the interaction of these regional astrocyte subtypes with other cell types. Figure 3d and Supplementary Figure 3d illustrate the interactions where ligands expressed by regional astrocyte subtypes activate other target genes in other cell types across samples S1 and S2 respectively. Figure 3e and Supplementary Figure 3e demonstrate the interactions where ligands expressed by other cell types activate specific marker genes in astrocyte subtypes. Ligands expressed by the telencephalic astrocytes in Cortex, and Amygdala were found to activate genes in excitatory neurons in Layer 6b (*Vegfa:Ctgf*) and Amygdala (*Ptn:Nr2f2*) respectively. On the other hand, *Agt* expressed by diencephalic astrocytes (Thalamus) were found to activate genes in inhibitory neurons (*Psap:Pde4dip*). White matter astrocytes interacted with oligodendrocytes via *Tnc* which activated gene *Adamts4*. Similarly, different genes in telencephalic astrocytes in Amygdala, Cortex, and Hippocampus were found to be activated by ligands expressed by excitatory neurons in Layer *4* (*Rspo1:Vegfa*), Amygdala (*Slit1:Nr2f2*), and Hippocampus DG (*Crlf1:Ccnd3, Slit1:Tcf4*) respectively. Excitatory neurons in the thalamus were found to activate genes in diencephalic astrocytes (*Ntng1:Gpld1, Nxph1:Tcf7l2*).

For each communication domain, based on the agreement of neighborhood activity and prior regulatory potential scores, we ranked the prominent ligands (Fig. 3f, Supplementary Fig. 3f) that are expressed by the major cell types in the domain (see Methods for details) which further highlighted the region-specific enrichment of certain ligands. For instance, *Bdnf* known to be associated with numerous functions within the hippocampus (39) displayed high activity within domain 16 associated with the hippocampus. *Nrg1*, known to have high impact on cortical functions (40) exhibited high activity within domain 5. Supplementary Table 2 lists such major ligands with their corresponding literature support. Using Renoir, we spatially mapped major pathway activities based on the activity scores of the ligand-target pairs associated with the pathways (see Methods) (Fig. 3g, Supplementary Fig. 3g and Supplementary Fig. 4). Consistent with prior works (41; 42), cortex and hippocampus were highly enriched in MAPK signalling and NEUROTROPHIN signalling pathways, respectively.

### Renoir uncovers intratumor spatial communication sub-domains within triple negative breast cancer

To demonstrate Renoir’s ability to characterize complex heterogeneous tissues in terms of ligand-target interactions, we analyzed single-cell and Visium datasets (Fig. 4a) from a triple negative breast cancer (TNBC) patient (28). Renoir quantified the spatial neighborhood activity scores for 18654 ligand-target pairs based on which, it clustered the spatial spots into six communication domains which in addition to being distinct in low-dimensional embeddings (Fig. 3b), were primarily associated with morphologically distinct pathological regions (Fig. 3c). The domains were further differentiated based on the ligand-target activities and the composition and abundance of the cell types they harbored (Fig. 4d-e, Supplementary Fig. 5a-b). Since two (clusters 4 and 5) of the six domains had low overall UMI count (Supplementary Fig. 5b), we focused on the first four communication domains in our further analyses. In accordance with their respective cell type compositions, interesting differential ligand-target activities were observed across the six domains. For instance, domain 0, histologically annotated as ductal carcinoma in situ (DCIS), harbored a high abundance of cancer cycling and cancer basal SC cell types (Fig. 4c, Supplementary Fig. 5a); cancer-associated fibroblasts (CAFs) and different T cell populations (Supplementary Fig. 5a); and demonstrated differential activity of MYC and SAA1 (*LAMB1-MYC, SERPINE2-MYC, SAA1-ESRRA*) known to be overexpressed in basal-subtype of breast cancer (43; 44; 45); and MDK (*MDK-CCNA2*), a known cancer progression marker (46). Domain 0 also harbored high activity of *VEGF* (*VEGFA, VEGFB*) (known to be highly up-regulated in breast cancer (47; 48; 49)), and other stromal ligands (*COL18A1, COL4A1*) (Fig. 4e). Domain 1 was associated with the histological regions lymphocytes and invasive cancer and majorly harbored cancer-associated myofibroblasts (myCAF), macrophages, and T cells (cycling and *CD8*^+^) along with the two cancer cell types (Fig. 4c, Supplementary Fig. 5a). Consequently, interactions associated with tumor progression and invasion (e.g., *CXCL10-MMP2, CCL3-ICAM1* (50), *FN1-MMP10* (51)), as well as suppression (e.g., *CXCL9-IRF1* (52)) among these cell types were enriched in domain 1 (Fig. 4e). Domain 2 was associated with the histological region normal, stroma and lymphocytes and harbored four major cell types including inflammatory-like CAFs (iCAF-like) and myoepithelial cells (Fig. 4c, Supplementary Fig. 5a). Interestingly, this domain harbored high activity of the ligand *CCL28* which is known to promote breast cancer and metastasis (53), target *HES1* which plays a crucial role in malignant cell proliferation (54) and *AREG* which is expressed by myoepithelial cells and loss of which can inhibit tumor proliferation, growth and invasiveness (55) (Fig. 4e). Domains 3 and 4, despite low UMI count, showcased distinct ligand-target activities (e.g., *IL6, IL33, IL1RN* in domain 3 and *PENK* in domain 4 (Fig. 4e)).

**Figure 4.**
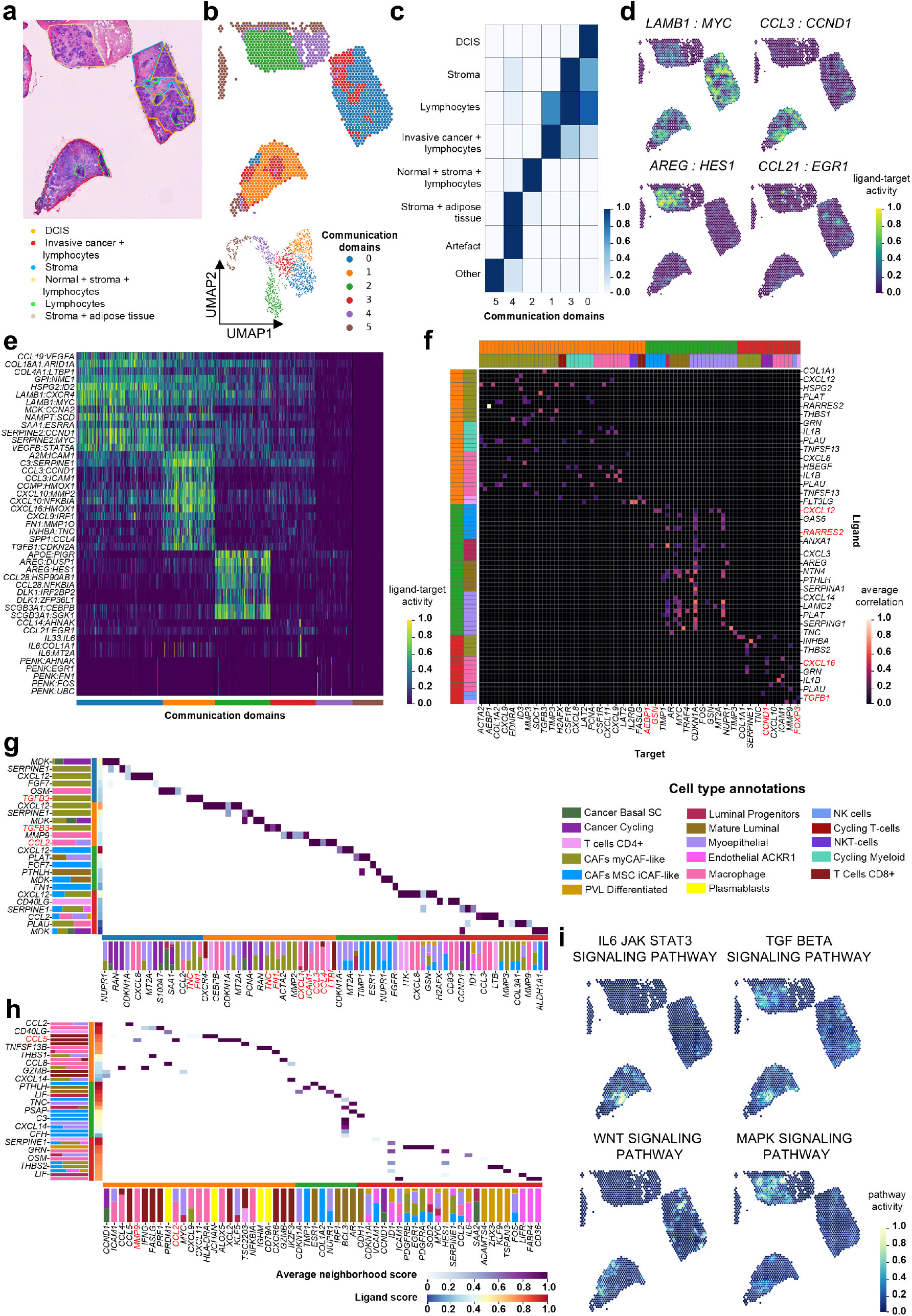
Renoir uncovers intratumor spatial communication sub-domains in triple negative breast cancer tissue. (a) The original histopathological annotations of H&E-stained triple negative breast cancer tissue (CID44971 (28), *n* = 1322 spots) showing six pathologically distinct regions. (b) Spatial communication domains inferred by Renoir based on ligand-target neighbor activity scores. The communication domains display distinct latent embeddings as can be seen from UMAP. (c) Distribution of spots in inferred communication domains across pathological annotations as in a. Each cell represents the proportion of spots of a communication domain in the corresponding pathological region. (d) Spatial map of neighborhood activity scores for the ligand-target pairs (e) Differentially active ligand-target pairs inferred by Renoir across all communication domains. (f) Cell type-specific ligand-target interactions for the major cell types in domains 1, 2, and 3. The inner color bar represents the cell types of the ligand (left) and target (top) and the outer color bar represents the communication domain. Each cell represents the average Pearson correlation between the ligand-target neighborhood scores and the abundances of the cell types expressing the ligand and target across the spots pertaining to the domain being considered. (g) - (h) Domain-specific ligand ranking based on their cumulative activities over target genes expressed by major cell types in the domain. (g) Top six ligands for communication domains 0, 1, 2, and 3 are represented. (h) represents the unique ligand-target pairs within each domain. Stacked color bars represent the cell types that express the ligand (right) and target (top). (i) Spatial map of four hallmark pathways - IL6 JAK STAT SIGNALING, TGF BETA SIGNALING, WNT SIGNALING and MAPK signalling pathway activity.

We further investigated the most prominent cell type-specific ligand-target interactions (Fig. 4f, Supplementary Fig. 6a) within each spatial communication domain (see Methods for details). By emphasizing domains with the highest ligand-target activities and UMI count (0, 1, 2, and 3), we uncovered cell type-specific interactions such as *RARRES2-AEBP1, CXCL12-GSN* (ligand and target are markers of iCAFs, domain 2), *CXCL16-CCND1* (domain 3). For each communication domain, we further ranked the major ligands (expressed by the major cell types in the domain) according to their activities (see Methods for details). A key EMT marker, *TGFB3* (56) was one of the top-ranked ligands in both domains 0 and 1 (which harbored a high abundance of cancer cells) and its top targets included *FN1* (in myCAFs and macrophages) and *TNC* (in myoepithelial cells and myCAFs) which are associated with breast cancer metastasis (57; 58), (Fig. 4g). In domain 1 which contained high abundance of macrophages, another top ligand was *CCL2*, which is known to promote metastasis via retention of metastasis-associated macrophages (MAMs) (59) and it was found to target *ICAM1* (a molecular target for TNBC) (50) in myoepithelial cells and macrophages, other chemokines *CCL3* and *CCL4* as well as lymphocyte inhibitory molecule *LTB* in *CD4*^+^ T cells (Fig. 4g). Within each communication domain, we also found ligands that were specifically active in a single communication domain (Fig. 4h). For example, in domain 1, *CD8*^+^ T cells secreted the ligand *CCL5* (known to promote TNBC progression through the regulation of immunosuppressive cells (60)) which activated *MMP9* and *CCL2* in macrophages.

Subsequently, utilising cell type abundance values of spatial locations, we sub-clustered each communication domain (domains 0, 1, 2, and 3) into subdomains that were unique in terms of cell type abundance and entailed distinct ligand-target activities (Supplementary Figs. 7-10). Amongst the six subdomains in domain 0 (Supplementary Fig. 7), in subdomain 4 *TGFB1* and *FBN1* activated several EMT-associated genes (*FBN1, MMP2, COL5A1, PDGFRB*, etc.) (61) and cancer progression and regulation markers (*DUSP1, JUNB*, and *MYC*) (62; 63) respectively. In contrast, subdomain 1 which harbored different immune cells (*CD4*^+^ and *CD8*^+^ T cells, DCs, NKT cells) as well as iCAFs, showcased extensive chemokine activity (*CCL5*) which targeted different genes in different immune cells and iCAFs (e.g., *EGR1*). Domain 1 was segregated into 4 subdomains (Supplementary Fig. 8), of which subdomain 2 abundant with immune cells showcased high *RARRES2* activity (secreted by myCAFs) and subsequently a lower abundance of cancer cells compared to other subdomains. This is coherent with previous findings which show that *RARRES2* serves as an immune-dependent tumor suppressor by acting as a chemo-attractant to recruit anti-cancer immune cells (64; 65; 66; 67). Similarly, subdomains of other domains showcased interesting activity particular to the cell type abundant across the subdomains (Supplementary Figs. 9-10).

Using Renoir, we further spatially mapped the major pathway activities based on the activity scores of the ligand-target pairs associated with the pathways (see Methods). While some pathways showed activity across multiple communication domains, some other pathways were more active in specific communication domains (Supplementary Fig. 6b). For example, the WNT signaling pathway was found to be active in subdomain 1 of domain 0, and subdomain 2 of domain 1 which harbored a high abundance of cancer cell types (Fig. 4i). IL-6/JAK/STAT3 signaling was highly active in subdomain 1 of domain 1 (Fig. 4i) that harbored a high abundance of myCAFs, macrophages and myoepithelial cells along with cancer cells but lacked immune cells other than cycling T-cells indicating a highly immunosuppressive tumour microenvironment (68). Taken together, Renoir was able to provide comprehensive insights into the spatial heterogeneity of ligand-target activity and its implication in tumor biology.

### Renoir identifies hepatocyte-macrophage interactions in developing fetal liver

Cell-cell communication shapes tissue functioning and organogenesis during development. In order to evaluate the versatility of Renoir in uncovering cell-cell interactions during development, we applied Renoir to data from fetal liver development. We generated a 10X Visium spatial transcriptomic dataset (see Methods for details) from a fetal liver sample for which single-cell RNA sequencing data was available (69) (Fig. 5a). Renoir identified 17630 ligand-target pairs active in the tissue, quantified their spatial neighborhood activity scores and based on these scores, identified five spatial communication domains (Fig. 5b), which differed based on ligand-target activity and distribution of UMI count across the tissue section (Fig. 5b-c and Supplementary Fig. 4a). Domain 0 demonstrated the majority of ligand-target activity, followed by domain 1. On the other hand, domains 2 and 4 showed sparse activity (UMI counts were low) (Fig. 5c) while domain 3 was specific to the target *CDKN1A* (Supplementary Fig. 4a). This is in line with the UMI count distribution across spots, where the UMI count was most abundant in domains 0 and 3, followed by domains 1, 2 and 4. Cell types were found to be evenly distributed across domains 0, 1 and 3 (Supplementary Fig. 11b). Activities pertinent to fetal liver were found to be widespread (Fig. 5d). For example, *COL18A1* associated with type XVIII collagen (a prominent component of the Extracellular Matrix (ECM) in the liver (70)) was found to be highly expressed across the tissue and activate multiple targets including hematopoietic transcription factors *BCL2L1* and *BCL6* (71). Higher *CDKN1A* activity was observed, in line with the proliferative nature of hepatocytes in developing fetal liver tissue.

**Figure 5.**
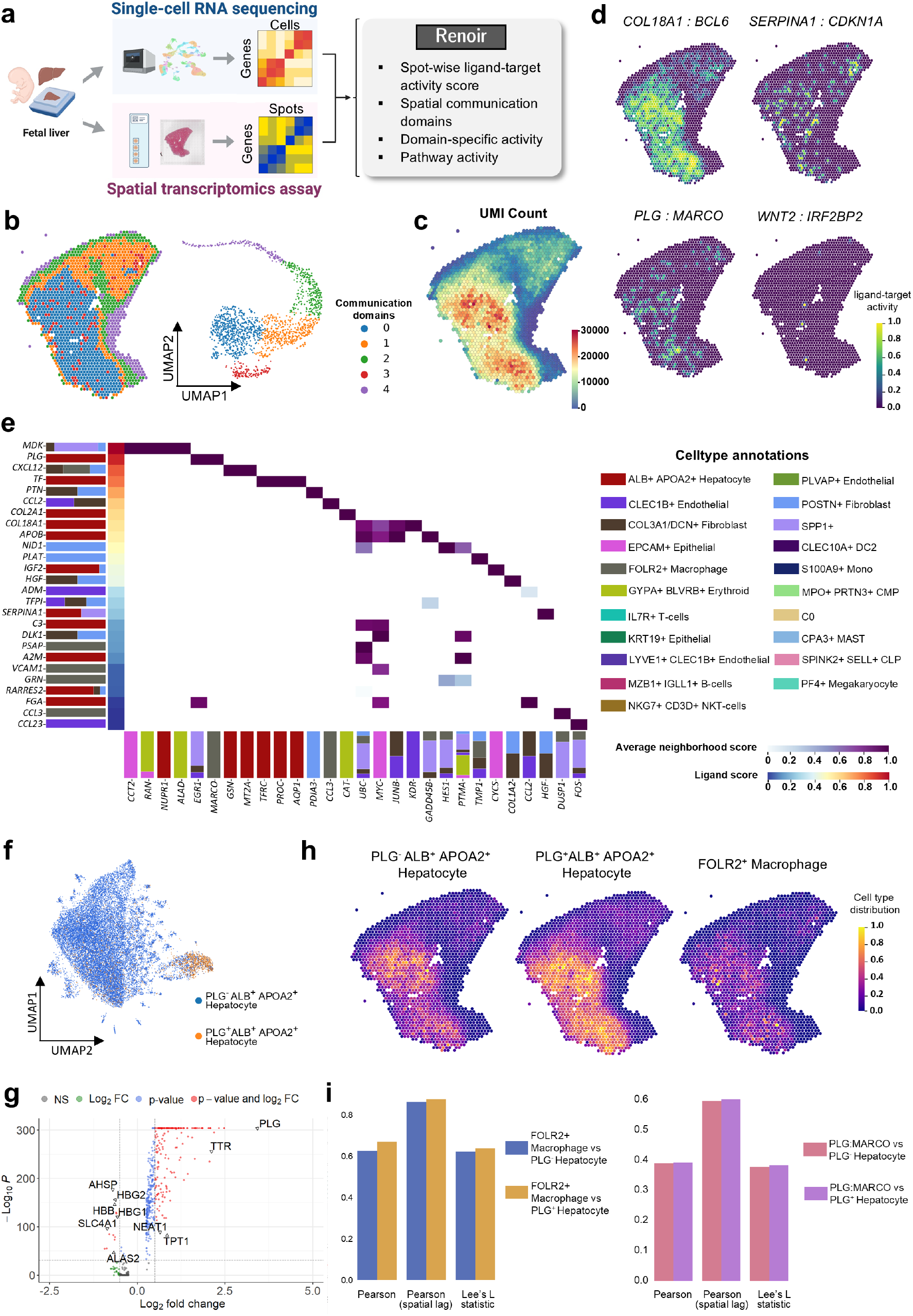
Renoir identifies hepatocyte-macrophage interactions in developing fetal liver. (a) Preparation of FFPE fetal liver tissue (b) Spatial communication domains inferred by Renoir based on ligand-target neighborhood scores. (c) Distribution of UMI count across the tissue section. (d) Spatial map of neighborhood activity scores for the ligand-target pairs (e) Ranking of ligands based on their cumulative activities over target genes expressed by major cell types in domain 0. (f) UMAP plot of latent embedding for hepatocyte population in scRNA-seq data, cells are colored as either *PLG*^+^ or *PLG*^−^. (g) Cell type abundances of Hepatocytes (*PLG*^+^ and *PLG*^−^) and *FOLR2*^+^ Macrophages overlaid onto the spots in ST data. (h) Volcano plot depicting differentially expressed genes between *PLG*^+^ and *PLG*^−^ Hepatocytes (−*log*_10_*P* threshold = 32 and *log*_2_*foldchange* threshold = 0.5). (i) Spatial similarity measures between *PLG:MARCO* neighborhood activity scores and cell type abundances of *PLG*^+^, *PLG*^−^ Hepatocytes and *FOLR2*^+^ Macrophages.

Further analyses involved acquiring previously described ligand rankings and prominent cell type-specific ligand-target interactions for each communication domain. Specifically, we observed prominent activity of the ligands *MDK*, an important growth factor for fetal liver hematopoietic stem cells and multipotent progenitor expansion (72) and *CXCL12* which helps in retaining hematopoietic stem cells in the fetal liver (73) (Fig. 5e). Importantly, we observed *MARCO*^+^ tissue-resident macrophage population (Kupffer cells) interacting with *APOB*^+^ hepatocytes. *MARCO* (MAcrophage Receptor with COllagenous structure) is predominantly expressed in non-inflammatory Kupffer cells (74) and fetal-like reprogramming of liver macrophages plays an important role in immunosuppressive phenotype in liver cancer (69). As shown in Fig. 5e and Supplementary Fig. 12a, we observed that *MARCO*^+^ macrophages interact with *PLG*^+^/*APOB*^+^ subset of hepatocytes. The *PLG* (plasminogen) has been known to be important for hepatic regeneration (75; 76) and *PLG* deficiency can lead to impaired remodelling in mouse models after acute injury. Next, we subdivided hepatocytes into two groups: *ALB*^+^*APOA*^+^*PLG*^+^ and *ALB*^+^*APOA*^+^*PLG*^−^ (Figs. 4f-g). As shown in Figs. 5g-h, these two subsets are transcriptionally and spatially distinct from each other indicating heterogeneity of the fetal liver hepatocyte population. We observed that the majority of the *ALB*^+^*APOA*^+^ hepatocyte population does not express *PLG* while there is a specific cluster with a high expression of *PLG* (Fig. 5f). In the Visium data, the spots expressing *ALB*^+^*APOA*^+^*PLG*^+^ hepatocytes also contained *FOLR2*^+^ macrophages (Fig. 5h). *FOLR2*^+^ macrophages are embryonic macrophages present in the fetal liver microenvironment (69). Spatial correlation analyses further showed higher similarity of *FOLR2*^+^ macrophages and *ALB*^+^*APOA*^+^*PLG*^+^ hepatocytes as compared to *PLG*^−^ hepatocytes in terms of cell type distribution as well as neighborhood activity score (Fig. 5i). Taken together, these results provide important insights into the macrophage-hepatocyte niche in the developing human liver.

### Renoir identifies onco-fetal interactions in hepatocellular carcinoma

To evaluate Renoir’s ability in handling single-cell resolution ST datasets, we applied it on a Nanostring CosMx dataset generated from a hepatocellular carcinoma (HCC) patient (77). The dataset consisted of 70166 cells distributed across 46 cell types and 959 genes.

We recently (69; 77) discovered the presence of onco-fetal tumor microenvironment (TME) in hepatocellular carcinoma, where the *FOLR2*^+^ TAMs, *PLVAP*^+^ ECs and *POSTN*^+^ CAFs were identified as onco-fetal cell types that shared transcriptional programs with their fetal liver counterparts. We evaluated Renoir’s performance in identifying the onco-fetal interactions from the single molecule imaging (CosMx) dataset. To perform an unbiased analysis, we used Renoir to quantify the activity of the ligands expressed by the onco-fetal cell types with the other major cell types present in the neighborhoods of the onco-fetal cells (Supplementary Fig. 13a) using Renoir’s ligand-ranking formulation (Fig. 6a). Renoir was able to characterize the activity of several known core onco-fetal ligands (e.g., *VEGFA, TGFB3*) as well as ligands shared by two onco-fetal cell types (e.g., *BMP7, CXCL12, TGFB1, IFNG*) (Supplementary Fig. 13b) (77). Specifically, Renoir identified the activity of known onco-fetal interactions *CXCL12* -*CXCR4* and *VEGFA*-*KDR* (Fig. 6c). *CXCL12-CXCR4* signaling was particularly elevated in onco-fetal niche where *CXCL12* was expressed by *PLVAP*^+^ ECs and *FOLR2*^+^ TAMs and it activated *CXCR4* in *POSTN*^+^ CAFs as well as macrophages and Tregs. Renoir correctly identified the co-localization of these cells and quantified the activity of the ligand-target pair in the neighborhood. Similarly, *VEGFA* was expressed by all three onco-fetal cell types and it interacted with *KDR* in co-localized ECs and *MUC6*^+^ bipotent hepatocytes (Fig. 6c).

**Figure 6.**
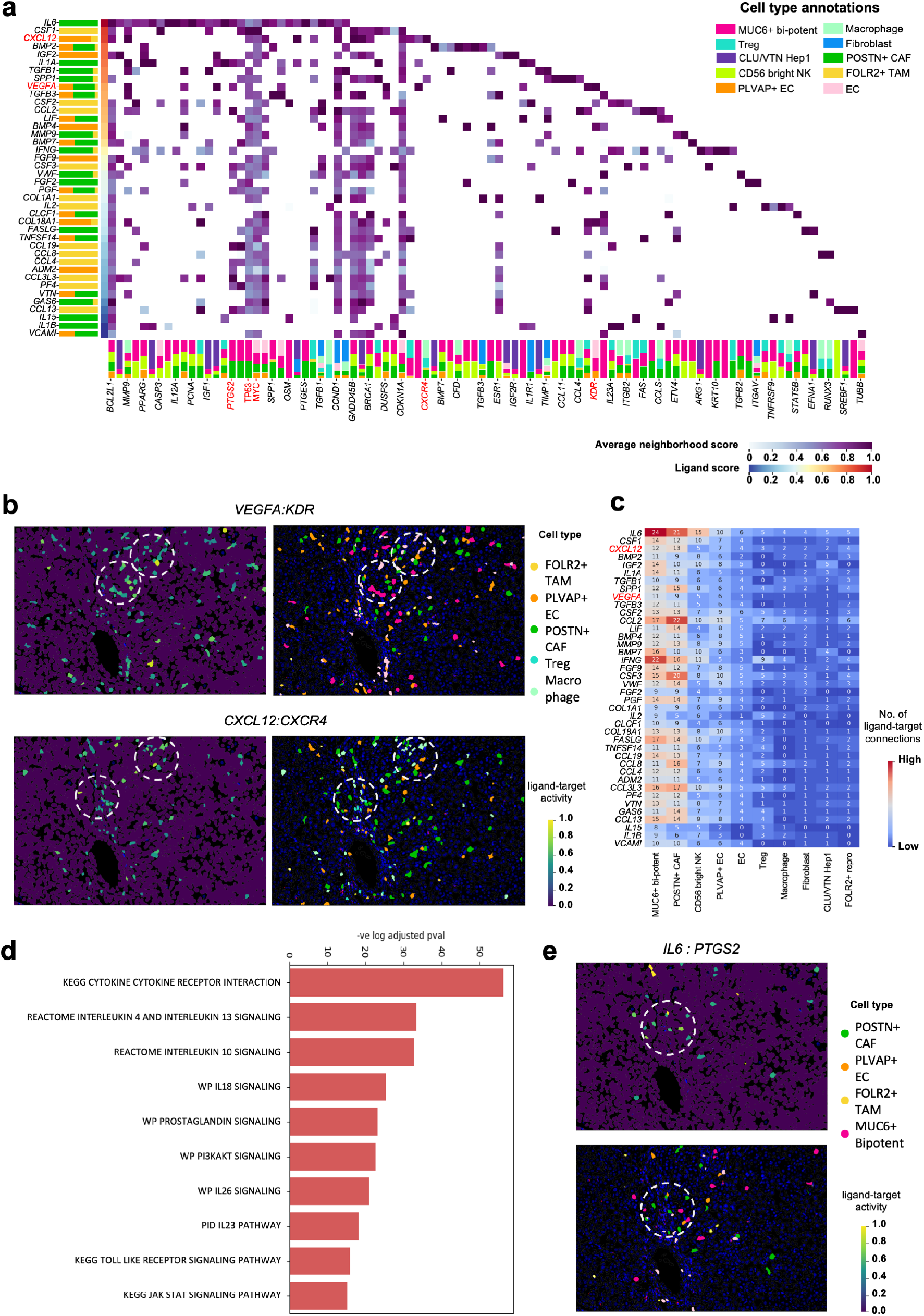
Renoir identifies onco-fetal interactions in hepatocellular carcinoma. (a) Ranking of activity of ligands expressed by onco-fetal cell types with other major cell types present in the neighborhoods of the onco-fetal cells. Stacked color bars represent the cell types that express the ligand (left) and target (bottom). (b) (left) Spatial map of cell type specific neighborhood activity scores across cell types associated with the ligand and target for *V EGFA* : *KDR* and *CXCL*12 : *CXCR*4 respectively (fov 10); (right) Spatial distribution of cell types associated with the ligand and target for *V EGFA* : *KDR* and *CXCL*12 : *CXCR*4 respectively (fov 10) (c) Distribution of ligand-target interactions across target cell types. The target cell types include major non-onco fetal cell types present in the neighborhoods of the onco-fetal cell types and onco-fetal cell types (d) Top pathways in the *MUC6*^+^ bi-potent cells targeted by the oncofetal ligands (e) (From left to right) Spatial map of cell type specific neighborhood activity scores across cell types associated with *IL*6 : *PTGS*2; Spatial distribution of cell types associated with *IL*6 : *PTGS*2.

### Renoir discovered onco-fetal niche mediated reprogramming of *MUC6*^+^ bipotent cells

Renoir not only validated known onco-fetal interactions but also identified novel biological interactions between onco-fetal niche and putative stem-like *MUC6*^+^ cells in liver cancer microenvironments. Renoir identified the *MUC6*^+^ bipotent hepatocytes to have the highest number of interactions with the onco-fetal cell types (more specifically with *POSTN*^+^ CAFs) (Fig. 6b). We delved into interactions specific to onco-fetal cells and *MUC6*^+^ bipotent cells by quantifying the cell type-specific activity of the relevant ligand-target pairs in the onco-fetal neighborhoods (Supplementary Fig. 13c). We noticed the activation of *MYC* and *TP53* in *MUC6*^+^ bipotent cells by multiple onco-fetal ligands including *BMP2, TGFB3*, and *CXCL12*. We utilized gprofiler (78) to infer the pathways in the *MUC6*^+^ bipotent cells targeted by the onco-fetal ligands and identified various interleukin signaling pathways (e.g., IL4, IL13, IL10), cytokine signaling and prostaglandin signaling (Fig. 6d) to be enriched. Since prostaglandin signaling is known to promote HCC progression (79), we further looked into the activity of *IL6-PTGS2*. High scores for *IL6-PTGS2* was found in the tissue niche where *MUC6*^+^ bipotent cells were co-localized with the oncofetal cells indicating the potential role of onco-fetal cells in the activation of PTGS2 in the *MUC6*^+^ bipotent cells (Fig. 6e). Importantly, these observations indicate that application of Renoir could lead to the identification of novel biological insights in the tumor microenvironment.

## Discussion

Cell-cell interactions play a crucial role in various biological processes, including tissue development, homeostasis, and disease. These interactions are essential for the proper functioning of multicellular organisms, as they facilitate communication and coordination among different cell types. Single-cell RNA-seq approaches have allowed us to comprehend the heterogeneity of cell types in a complex tissue. However, traditional single-cell RNA-seq data lacks spatial context, making it challenging to uncover the spatial organization and intercellular relationships that underpin complex biological systems. Spatial transcriptomics is a burgeoning field that addresses this limitation by providing spatially resolved gene expression data, enabling researchers to explore cells in a spatial context and bridging the gap between molecular and morphological observations.

In this regard, our newly developed computational method, Renoir, offers a powerful tool for investigating cellular interactions within their native spatial context, from development to disease. Renoir enables the spatial mapping of ligand-target activities through the quantification of a neighborhood activity score, infers spatial communication domains by grouping spots based on the similarity of ligand-target activities and uncovers ligand-target interactions restricted to each communication domain. Renoir further enables the identification of the major ligands active in each domain and the spatial mapping of pathway activities based on known or denovo genesets. Renoir can work on datasets generated using spot-resolution spatial technologies as well as singlecell resolution spatial technologies.

Our comprehensive benchmarking of Renoir on a variety of datasets simulated from real datasets across tissue types demonstrate its superiority over existing CCC inference methods in quantifying the ligand-target activities across spatial locations. Specifically, by accounting for the receptor expression in the target cell type, Renoir is capable of reducing false positive interactions which are abundantly found by other methods. Using DLPFC spatial datasets with ground truth annotations of spatial domains, we further illustrated that the communication domains inferred by Renoir can better capture the underlying spatial structure of the tissue as compared to other methods.

Our application of Renoir to three distinct 10X Visium spatial transcriptomics datasets including adult mouse brain, human fetal liver, and human triple negative breast cancer (TNBC) demonstrate Renoir’s versatility in characterizing ligand-target interactions across tissue types. For each dataset, Renoir identified distinct spatial communication domains which harbored distinct ligand-target activities pertaining to the major cell types abundant in the domain and these were also consistent with known literature. In the adult mouse brain, Renoir characterized the interactions of different regional astrocyte subtypes with other co-localized excitatory and inhibitory neurons. For the triple negative breast cancer, Renoir uncovered spatial communication domains and subdomains harboring distinct chemokine interactions between cancer, stromal and immune cells which were associated with tumor proliferation and metastasis. Renoir further identified signaling pathways that were active in cancer (WNT signaling) or immune/stromal cell type (IL-6/JAK/STAT3 signaling) dominated domains of the TNBC tissue. For developing fetal liver, Renoir uncovered an interaction between hepatocytes and macrophages mediated via the ligand *PLG*. The application of Renoir on a single-molecule imaging (Nanostring CosMx) hepatocellular carcinoma dataset further shows Renoir’s utility in handling single-cell resolution large-scale spatial datasets. From the hepatocellular carcinoma dataset, Renoir identified several onco-fetal interactions involving endothelial cells, macrophages and CAFs and uncovered the role of these onco-fetal cell types in activating prostaglandin signaling in the bipotent hepatocytes.

By integrating single-cell RNA-seq and spatial transcriptomics data, Renoir enables a more comprehensive understanding of the dynamic relationships between different spatially co-localized cell types and their roles in various biological processes. This approach allows for the identification of key players in cellular communication and provides insights into the mechanisms governing tissue organization, functionality, and response to disease. The several visualization modules provided by Renoir further enable the user to explore the spatial map of ligand-target and pathway activities of their interest. Ultimately, Renoir has the potential to advance our understanding of complex biological systems and contribute to the development of novel therapeutic strategies targeting cellular interactions.

## Methods

### Overview of Renoir

Using either paired cell type annotated scRNA-seq data and spatial transcriptomic (ST) data or single-cell resolution spatial transcriptomic data, and a set of curated ligand-target pairs, Renoir calculates spatially enriched neighbourhood scores for the curated set of ligand-target pairs for a given sample. The neighbourhood scores can subsequently be leveraged to track ligand-target activity across the geographic topology, as well as identify spatially enriched communication regions, cell type-specific interactions, and pathway activity.

### Required input data

#### Single-cell RNA-seq and Spatial transcriptomic data

As input, Renoir requires the following - (1) a spatial transcriptomics dataset *X*^*st*^ ∈ ℝ^*S*×*G*^ with gene expression for *S* spots and *G* genes. Depending on the technology, the spots can contain a mixture of cells (e.g., 10X Visium) or single-cell (e.g., Nanostring CosMX). In case of cell-resolution ST data, cell type annotations of the spots to one of *C* cell types is required. (2) Spatial coordinates for each spot, and (3) a corresponding cell type annotated single-cell gene expression dataset 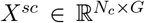 for *N*_*c*_ cells and *G* genes (required for ST datasets where each spot contains mixture of cells), where each cell mapped to one of *C* cell types. Note that the single-cell gene expression dataset is optional when single-cell resolution spatial transciptomic dataset is available.

#### Acquiring curated ligand-target pairs

We first curate a set of secreted ligands from NATMI’s connectomeDB2020 (8) database. For each ligand, we obtain a list of putative targets sorted according to the descending order of ligand–target regulatory potential scores as calculated by NicheNet’s prior knowledge model (11). The number of targets per ligand is user-defined. For our analyses, we have used the top 100 targets for each ligand. The ligand-target pair is retained for further analysis if both genes are among the highly variable genes in the spatial transcriptomic dataset. Thus, we obtain a set of ligand-target pairs, 𝒮 _*lt*_ specific to our spatial transcriptomic dataset.

### Calculation of neighborhood activity score for a ligand-target pair for a spot

#### Constructing neighborhoods

We utilize the spatial transcriptomic data to construct a directed graph such that each spot within the dataset represents a node in the graph and there exists a directed edge between a node (spot) and every node within its neighborhood. A neighborhood for a spot is defined with respect to the distribution of spots within the spatial dataset considered.

For data wherein the spots are equidistant from their immediate neighbors (e.g., 10X Visium and Spatial Transcriptomics), the neighborhood for a spot *s* is defined to be a set of spots inclusive of *s* such that every spot other than *s* is at a “one-hop” distance away from *s*. For data wherein the spots are not uniformly distributed across the sample (e.g., Slide-seq, StereoSeq, CosMx), the neighborhood for a spot *s* is defined to be a set of spots (inclusive of *s*) which are at most *r* unit distance away from *s*, where *r* is user defined.

#### Formulating neighborhood activity scores

On constructing the neighborhood graph, we quantify the potential interaction between a ligand *l* and target *t* within the neighborhood of a spot *s* using a neighborhood activity score defined by,

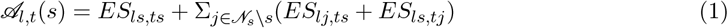

where 𝒩_*s*_ denotes the neighborhood of *s* and 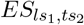 denotes the edge score (edge weight) of an edge directed from a spot *s*_1_ expressing ligand *l* to a neighboring spot *s*_2_ expressing target *t* and is defined by,

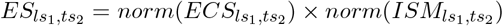

where 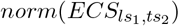 and 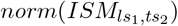 are the min-max normalized expression correspondence score and inherent similarity measure for the ligand-target pair *l* − *t* for the spots *s*_1_ and *s*_2_ respectively. The edge score quantifies the possible interaction between a ligand *l* at spot *s*_1_ and target *t* at spot *s*_2_ as the product of the inferred similarity between the expressions of *l* and *t* across all the cell types present at *s*1 and *s*2 respectively 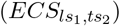 and the inherent similarity (acquired from the single cell data) between the expressions of *l* and *t* amongst cell types present at *s*_1_ and *s*_2_ respectively.

#### Calculation of Expression Correspondence Score and Inherent Similarity

##### Measure

To calculate the Expression Correspondence Score and Inherent Similarity Measure between two neighboring spots *s*_1_ and *s*_2_ where a ligand *l* is expressed at spot *s*_1_ and target *t* is expressed at spot *s*_2_, we utilize the following formulations

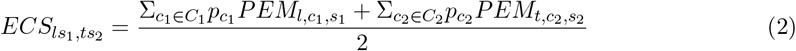

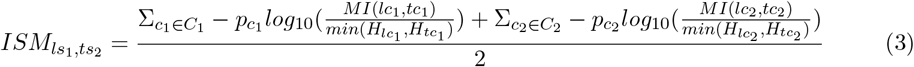

where, *C*_1_ and *C*_2_ denote the set of cell types present at *s*_1_ and *s*_2_ respectively, 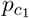 and 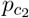 denote the cell type proportions for cell types *c*_1_ ∈ *C*_1_ and *c*_2_ ∈ *C*_2_ and *c*_2_ expresses at least one receptor of *l* respectively. *c*_2_ is considered to express a receptor if the receptor gene is expressed in more than *thresh*% of cells of that cell type (computed based on scRNA-seq/single-cell ST data), where *thresh* in our analysis was set to 5 following NicheNet. 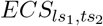 is defined as the average of cell type proportion-weighted PEM of ligand *l* and target *t* in spots *s*_1_ and *s*_2_ respectively. The PEM value for cell type specificity of gene *g* in cell type *c* for spot *s* is given by

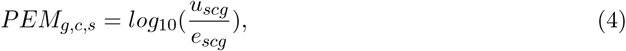

where, *u*_*scg*_ and *e*_*scg*_ denote the inferred and expected expression of gene *g* specific to cell type *c* at spot *s* respectively. In eq. (3), *MI*(*lc*_1_, *tc*_2_) denotes the mutual information between the expression of *l* across cell type *c*_1_ and the expression of *t* across cell type *c*_2_. *H*(*lc*_1_) denotes the Shannon entropy of the expression of ligand *l* across cell type *c*_1_ and *H*(*tc*_2_) denotes the Shannon entropy of the expression of target *t* across cell type *c*_2_. *MI*(*lc*_1_, *tc*_2_), *H*(*lc*_1_), and *H*(*tc*_2_) have been estimated from the single-cell (either scRNA-seq or single-cell ST) data using the Miller-Madow correction.

#### Quantifying inferred and expected cell type-specific gene expression

In order to quantify the cell type-specific gene expressions at each spot, we estimate the abundance of all cell types in each spot (*W*_*S*×*C*_) and gene-specific scaling parameters (*M*_*G*×1_) using Cell2location (31) that takes as input the ST data *X*^*st*^ and reference cell type signatures (𝒢^*C*^ : *C* × *G*) estimated from the single-cell data *X*^*sc*^. We then compute the absolute number of mRNA molecules contributed by each cell type *c* to each spot *s* to quantify the gene expression of a specific gene *g* at spot *s* for cell type *c* as given by:

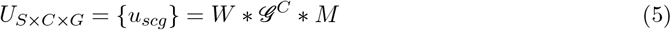

where * represents matrix multiplication without sum-reduction. For single-cell resolution datasets, the cell type annotation of the spots are directly used as cell type abundance information, where *W*_*S*×*C*_ represents the cell type annotation of each spot and *U*_*S*×*C*×*G*_ represents the original single-cell resolution ST dataset. This gives us,

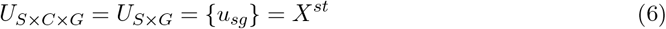

The expected expression of a specific gene *g* at spot *s* for cell type *c* can then be calculated as:

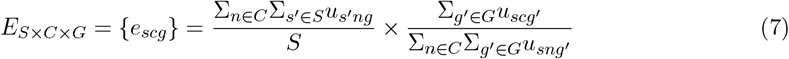

For single cell resolution datasets, since each cell is of a specific cell type the above formulation collapses to,

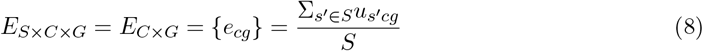

#### Inference of spatial communication domains based on ligand-target neighborhood scores

In order to identify distinct spatial communication domains, which we define as a group of spots harboring similar ligand-target interactions, we performed clustering of the spots based on the ligand-target neighborhood activity scores. After computing the ligand-target neighborhood scores for all curated ligand-target pairs, each spot *s* can be represented as a vector of ligand-target activity scores, 𝒜_*l*,*t*_ of ligand-target pairs (*l, t*) ∈ 𝒮 _*lt*_ and based on these ligand-target activity score vectors, spots are clustered using the scanpy library (80). First, the top 2000 highly variable ligand-target pairs are identified followed by principal component analysis. Leiden clustering is performed on the k-nearest neighbor graph obtained based on the principal components.

#### Inferring cell type-specific sub-communication domains

Additionally, we further subclustered the spots present within each of the spatial communication domains based on the cell type abundance of the spots. For sub-clustering also, Leiden clustering was adopted and implemented using scanpy.

### Inference of communication domain-specific ligand-target activities

#### Differential ligand-target activity across communication domains

The Wilcoxon-rank sum test was utilized to estimate the ranked, differentially active ligand-target pairs for each communication domain using scanpy’s ‘rank genes groups()’ function.

#### Ranking of ligands based on activity within communication domains

Renoir further allows the ranking of ligand activity within a communication domain. In order to perform ranking of ligand activity, we first determine the major cell types (cell types that together constitute ≥ 90% of cell type abundance) within a communication domain based on the average proportion of the cell types across all the spots within that domain. For each major cell type in a domain, we identify the top marker genes (*n* = 500) from the scRNA-seq dataset. We acquire a list of ligand-receptor pairs from NATMI’s connectomeDB (8) and identify the major ligands that are also present within the marker gene lists of the major cell types within the domain. For each ligand, we identify a set of marker targets across the major cell types within the domain. A target is selected if it falls within the marker gene list of the cell type and the cell type expresses a receptor for that ligand. For each ligand, we compute the average neighborhood activity score for each of its marker targets (averaged across the spots expressing the associated receptor cell type). Each ligand is then scored based on the Pearson correlation between the average neighborhood scores of the ligand-target pairs of the cell type marker targets and the regulatory potential scores for each ligand-target pair acquired from NicheNet’s prior model (11). Using the ligand score, Renoir ranks the ligands, where the ranking represents the amount of agreement between the ligand-target activity in our dataset and the prior knowledge of the pairs. While in our analyses, we used ligand-receptor and ligand-target pairs from specific sources, Renoir can also work with user-defined ligand-receptor and ligand-target pairs.

#### Inferring major cell type-specific ligand-target activity within a communication domain

To identify cell type-specific ligand-target activity within each communication domain, we first considered the ligand-target pairs for which both the ligand and target fell within the top 200 markers for the major cell types of the communication domain. For each domain where the ligand and target were markers of the major cell types, a simple average of the Pearson correlations between the neighborhood activity scores of the ligand-target pair and the cell type proportions of the cell types expressing the ligand and target respectively were calculated across the spots pertaining to the domain being considered. The correlation score measures how prominent a cell type-specific ligand-target activity is within a communication domain.

### Inference of spatial mapping of pathway activity

#### Construction of genesets of known pathways or Denovo clusters

Renoir further allows the spatial mapping of a pathway activity by utilizing existing or denovo gene sets. First, for a particular pathway represented by a gene set, we determine the ligand-target pairs active in the dataset by intersecting the set of all possible gene pairs within a gene set with the set of curated ligand-target pairs. In this work, we restricted our usage to the GSEA Molecular Signatures Database (81; 82), specifically HALLMARK and CANONICAL PATHWAYS (KEGG, WIKIPATHWAYS) gene sets. Renoir can also utilize any predefined gene set provided by the user, specific to the user’s requirements. Apart from utilizing gene sets for existing pathways, Renoir can also identify de novo clusters of ligand-target pairs based on their similarity of the neighborhood activity scores across the spots. For performing the clustering of ligand-target pairs, Renoir employs hierarchical clustering strategies, namely HDBSCAN and dynamic hclust. HDBSCAN was implemented using the hdbscan python package with the min pts parameter set to the lowest possible value and dynamic hclust was implemented using the dynamicTreeCut package.

#### Spatial mapping of existing pathways and Denovo clusters

To generate a spatial map of previously known pathways or Denovo clusters, we computed the first principal component after performing dimensionality reduction over the neighborhood activity scores of the subset of ligand-target pairs corresponding to a pathway/Denovo cluster using the ‘TruncatedSVD’ function of sklearn. This generates a pathway activity score for each spot in the spatial transcriptomics data.

### Benchmarking of Renoir

#### Generation of simulated data

For benchmarking the performance of Renoir, we simulated spatial datasets based on a semisynthetic approach inspired from the simulation approach in (83). In order to simulate a dataset with known spatial interaction between ligand and target, we start with a paired scRNA-seq and ST data (the reference dataset) and cell type deconvolution of the reference ST dataset is performed using cell2location (31). After that, we simulate the ST dataset using the following steps in order to maintain a controlled environment, where we can keep track of the different kinds of interactions enriched within the data.

1. Select a set of ligand-target pairs *LT* such that both the ligand and target are marker genes (top 100 markers) of at least one cell type in the paired scRNA-seq data. After that, the pairs in *LT* are segregated into two mutually exclusive subsets: *LT*_*m*_ and *LT*_*s*_. *LT*_*m*_ consists of the ligand-target pairs where the target gene is a marker for multiple (two or more) cell types and *LT*_*s*_ consists of every other pair, where the target gene is a marker of exactly one cell type. In both *LT*_*m*_ and *LT*_*s*_, the ligand could be a marker of one or more cell types. In our analysis, we were able to find a maximum of two cell types associated with each of the ligand or target genes.
2. For a specific ligand-target pair *lt* ∈ *LT*_*s*_, where the target is a marker of a cell type *ct*_*t*1_; in the ST dataset, we generate two types of reference spots, positive and negative spots respectively. Positive reference spots denote the spots within the tissue where ligand-target interactions for *lt* are simulated through the enrichment of the ligand cell type and target cell type *ct*_*t*1_, where a spot is enriched with a cell type by adding *n*_*e*_ = *Uniform*(2, 5) single cells of the said cell type into said spot. The UMI counts are then scaled back to the original UMI count to maintain the original UMI distribution of the sample being simulated. Negative references refer to the spots within the tissue that do not harbor any activity for *lt* as indicated by the low proportion of ligand cell type and target cell type *ct*_*t*1_. Specifically, negative reference spots are selected as a set of spots for which both ligand cell type and target cell type *ct*_*t*1_ proportions are low and their neighbors also have low proportions (below acceptable threshold) of ligand and target cell types. In the negative reference spots, interactions of *lt* can not occur given the low abundance of both the ligand and target cell types in these spots as well as in the neighboring spots.
3. For a specific ligand-target pair *lt* ∈ *LT*_*m*_ where the target is a marker of two cell types *ct*_*t*1_ and *ct*_*t*2_; in the corresponding ST dataset, we generate two types of reference spots, positive and negative. Without loss of generality, we assume that the cell type *ct*_*t*1_ does not express a known receptor of the ligand *l* but the cell type *ct*_*t*2_ expresses a known receptor of *l* (according to a ligand-receptor database, e.g., NATMI). Positive references are those spots within the tissue that have been enriched with the ligand cell type and target cell type *ct*_*t*2_. Specifically, a set of spots are selected that are not devoid of the ligand cell type. This is to ensure that the enrichment follows the underlying distribution of cell types in the dataset. For each of the selected spots, the spot is enriched with the ligand cell type and the neighbors of the spot are enriched with the target cell type *ct*_*t*2_. Negative references are those spots within the tissue that have been enriched with the ligand cell type and target cell type *ct*_*t*1_ (type 1 negative); or spots within the tissue that do not harbor any activity for *lt* as indicated by the low proportion of ligand cell type and target cell type *ct*_*t*2_ (type 2 negative). To simulate type 1 negative spots, a set of spots are selected from the set of all spots with low *ct*_*t*1_ and *ct*_*t*2_ proportions. We then enrich these spots with the ligand cell type and the target cell type *ct*_*t*1_. Thus for the type 1 negative spots the ligand-target interactions pertinent to the ligand cell type and target cell type *ct*_*t*1_ could not occur since no known receptors are expressed by *ct*_*t*1_ despite the co-localization of the cell types. The simulation of type 2 negative spots follows the same strategy as in case of *lt* ∈ *LT*_*s*_ considering *ct*_*t*2_ as the target cell type.

#### Scoring and evaluation strategy

To evaluate the performance of a CCC method in inferring the ligand-target activity at spot resolution, we employ the following scores.

#### Accuracy of spatial activity

The true activity of a ligand-target pair across the positive and negative reference spots is computed by employing the local Moran’s bivariate test-statistic 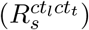 between the ligand and target cell types,

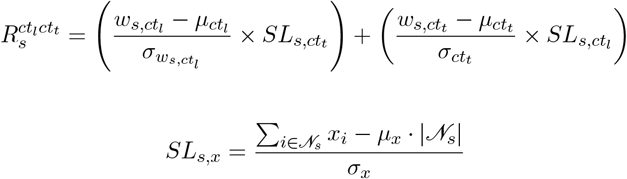

where *w*_*s*,*x*_ represents the cell type abundance of cell type *x* at spot *s, µ*_*x*_ represents the mean cell type abundance of cell type *x* across all spots considered, *σ*_*x*_ represents the standard deviation of cell type abundance of cell type *x* across all spots considered and *SL*_*s*,*x*_ represents the spatial lag of cell type *x* at spot *s*. This provides a measure of local interactions between the ligand and target cell types. Pearson’s correlation (as discussed before) is then used to calculate the average correlation between the ligand-target activity scores inferred by each method and 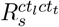 which gives a measure of how well does the inferred activity scores align with the true interactions as quantified in terms of spatial cell type co-localization.

#### Precision and Recall

To calculate the precision and recall of a CCC inference method in inferring the spatial interactions of ligand and target, we divide the spots into two sets based on a predefined threshold set on the ligand-target scores computed by the CCC inference method. Given a specific method, the spots which have a score greater than or equal to the threshold are considered to be positive predicted spots (harboring the ligand-target interaction) and the spots with a score lesser than the threshold are considered to be negative predicted spots (no ligand-target interaction). We then consider the these positive and negative predicted spots and compare them against the ground truth positive and negative reference spots. A predicted spot is considered to be a true positive (TP) if the spot is a positive predicted spot and is also a positive reference spot, true negative (TN) if the spot is a negative predicted spot and is also a negative reference spot, false positive (FP) if the spot is a positive predicted spot but is a negative reference spot and a false negative (FN) if the spot is a negative predicted spot but a positive reference spot. For each method, we calculate the TP, TN, FP and FN values to calculate the precision and recall of a method for a given simulated dataset. Thus precision and recall are calculated for each simulated ligand-target pair and the final precision and recall for a dataset for a specific threshold is computed to be the average of the precision and recall values across all considered ligand-target pairs for the same dataset. Instead of selecting a single threshold, the threshold is varied to determine the precision-recall curve for each method across various threshold values.

#### Comparison of communication domains with ground truth layers in DLPFC dataset

In order to estimate how coherent the inferred communication domains are with known regions of the dataset, we first estimate the communication domains across all 12 DLPFC samples for each of the methods (Renoir, COMMOT, SpatialDM and stLearn). This is done so by applying leiden clustering (resolutions range from 0.1 to 1.6 in intervals of 0.1) over the ligand-target interaction scores generated by each of the methods across all samples. The NMI and ARI values are calculated between the cluster labels and ground truth annotations; for each of the methods across all samples over all resolutions. At each resolution, the averaged NMI and ARI values are calculated for each method across all samples. For each method, the cluster labels corresponding to the resolution at which the averaged NMI and ARI values is the best for said method in each sample, is now considered as the best communication domain inferred by a method for a specific sample. This results with a list of best NMI/ARI values for each method across each sample. The box plots are then generated over the final inferred NMI/ARI values.

#### Preparation of fetal liver data

FFPE fetal liver tissue was obtained under ethical approval from the Centralised Institutional Research Board of the Singapore Health Services, Singapore SingHealth and National Health Care Group Research Ethics Committees. To determine the quality of RNA in FFPE tissue, RNA was extracted from 10*µm* sections using a Qiagen FFPE RNA extraction kit (#73504) and following the manufacturer’s protocol. RNA quality was determined as “good” if the percentage of RNA fragments *>*200 nucleotides (DV200%) was at least 50% of all fragments. Next, a 5*µm* tissue section was placed on spatially barcode oligonucleotides, within the capture area (6.5 x 6.5*mm*) of the Visium slide (4 capture areas/slide) and processed according to the manufacturer’s protocol. Briefly, Visium slides with affixed FFPE tissue sections were deparaffinised in fresh xylene and rehydrated in reducing ethanol (in water) concentration gradient. The slides were stained with Haematoxylin and Alcoholic Eosin and imaged at 10 times optical magnification using a Nikon Ni-E Microscope. RNA was then de-crosslinked and tissue sections were hybridised with Visium Human Transcriptome Probe Set, overnight at 50°C. The following day, the slides were washed with supplied post-hybridisation wash buffer and hybridized probe pairs were ligated using supplied ligase buffer at 37°C. After ligation, the slides were washed with room temperature and pre-heated post-ligation wash buffer. The ligated probes were then released by digesting hybridized RNA with supplied RNase enzyme at 37°C. The tissue sections were then permeabilised at 37°C using the supplied permeabilising enzyme. Permeabilization enabled ligated probes to be captured by spatially barcoded oligonucleotides immobilised on the Visium slide. Captured hybridised probes were then extended using probe extension mix at 45°C and extended probes were eluted accordingly. To determine appropriate amplification cycle numbers for library preparation in the next step, 1*µl* of eluted probes were quantitively amplified (qPCR) in accordance with the manufacturer’s instructions. Quantification cycle (Cq) value at 25% peak relative fluorescence unit (RFU) was used to set cycle numbers during library preparation. The library was then quantified using TapeStation (Agilent High Sensitivity D5000) to determine the average fragment size and concentration before sequencing.

#### Integration of mouse brain datasets

The two mouse brain samples were integrated using the stLearn workflow (22). Neighborhood scores generated for each mouse brain sample were concatenated and min-max normalized across ligand-target pairs for every spot. The principal components calculated over the spatial transcriptomic data of each sample via scanpy were used as input to harmony algorithm (84), which projected the spots onto a shared embedding space and thus presenting a new set of adjusted principal components for the concatenated dataset. Leiden clustering was performed via scanpy over the concatenated dataset with adjusted principal components at a resolution of 1.3 to obtain a set of spatial communication domains shared across the two brain samples.

#### Mapping of regional astrocyte subtype activity in mouse brain

In order to find localized astrocyte subtype activity in appropriate regions of the mouse brain dataset, we first identified the top 100 marker genes for every cell type (including astrocyte sub-types) using the available mouse brain snRNA-seq dataset. We then identified the top 100 differentially active ligand-target pairs for each region of the Mouse Brain dataset. Two sets of pairs were then generated, (i) the first set consisted of ligand-target pairs where the ligand was a marker gene of a specific astrocyte subtype, the ligand-target pair displayed differential activity in a region associated with said astrocyte subtype and the target was a marker of a cell type associated with said region (ii) the second set consisted of ligand-target pairs where the target was a marker gene of a specific astrocyte subtype, the ligand-target pair was differentially active in a region associated with said astrocyte subtype and the ligand was a marker of another cell type associated with said region. The above steps were carried out using the scanpy library ‘rank genes group’ function.

#### Correlation measures used for analyzing *PLG:MARCO* interaction in fetal liver

The following correlation measures were utilized to measure the similarity between *PLG:MARCO* neighborhood scores with *PLG*^+^ and *PLG*^−^ hepatocytes and *FOLR2*^+^ Macrophages for the Fetal Liver dataset,

- *Pearson correlation*: We utilized Pearson correlation to find the correlation amongst specific ligand/target expression values (either derived from scRNA-seq data or spatial transcriptomic data) and cell type abundances. We also used Pearson correlation to acquire an initial estimate of the correlation amongst the neighborhood scores of a specific ligand-target pair with the abundances of the cell types expressing the ligand and the target. The overall correlation measure can be summarised as,

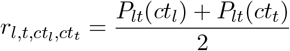

where,

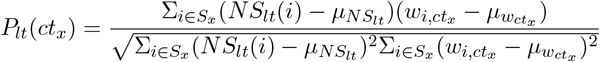

*l* and *t* represent a specific ligand and target respectively *ct*_*l*_ and *ct*_*t*_ represent the cell types expressing the ligand and the target respectively (as a marker) *NS*_*lt*_(*s*) represents the neighborhood scores of ligand-target pair *lt* at spot *s NS*_*lt*_ represents a vector of neighborhood scores of ligand-target pair *lt* across all *S*_*x*_ spots *w*_*s*,*y*_ represents the cell type abundance of the cell type the ligand(*ct*_*l*_) or target(*ct*_*t*_) is a marker of at spot *s* 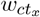 is a vector of cell type abundances of the cell type the ligand(*ct*_*l*_) or target(*ct*_*t*_) is a marker of, across all *S*_*x*_ spots 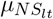 is the average neighborhood score for a specific ligand-target pair *lt* 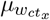 is the average cell type abundance for a specific cell type *ct_x_* 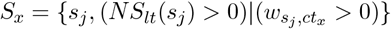 where *x* is either a ligand *l* or target *t*.

- *Pearson correlation over spatial lag* : We furher used Pearson correlation over spatial lag as a measure of spatial correlation (85). Pearson correlation over spatial lag considers the sum of expression or abundance values over a spot and its corresponding neighbors before applying Pearson correlation. Formally, Pearson correlation in the context of expression or abundance values can be defined as,

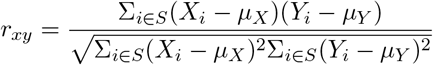

*X* and *Y* are the spatial lags defined by,

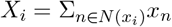

where *N* (*x*_*i*_) represents the neighborhood of 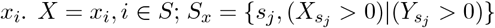 The spatial lags are calculated with respect to gene expressions from the spatial transcriptomics data or the spatial map of cell type abundances (*x* or *y*). While considering neighborhood scores, the scores are considered as they are since the neighborhood score at each spot takes into account the neighbors of the spot.

- *Lee’s L statistic*: We utilised Lee’s L statistic bivariate spatial association measure to incorporate spatial autocorrelation alongside spatial patterning between the two variables of interest (85). In our case, we adopted the measure to view the spatial co-patterning of neighborhood scores, gene expressions from the spatial transcriptomics data or the spatial map of cell type abundances with one another. This is defined as,

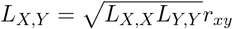

where,

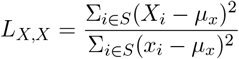

## Data availability

The original public data that was utilised in this study are provided below: snRNA-seq from adjacent sections in the mouse brain: downloaded from ArrayExpress under accession number E-MTAB-11115; Visium from adjacent sections in the mouse brain: downloaded from ArrayExpress under accession number E-MTAB-11114 (sample S1 = ST8059048 and sample S2 = ST8059052); Triple Negative Breast Cancer spatially resolved transcriptomics data: downloaded from the Zenodo data repository (https://doi.org/10.5281/zenodo.4739739); Triple Negative Breast Cancer raw scRNA-seq data: Downloaded from the Gene Expression Omnibus under accession number GSE176078. The scRNA-seq data for fetal liver was obtained from the Gene Expression Omnibus under accession number GSE156337. 10X Visium data for the fetal liver generated for this manuscript is available here. The Nanostring CosMx data for hepatocellular carcinoma has been obtained from https://doi.org/10.6084/m9.figshare.23972991.

## Code availability

Renoir has been implemented in Python and is freely available at https://github.com/Zafar-Lab/Renoir under the MIT license.

## Acknowledgements

This work was funded by the DBT/Wellcome Trust India Alliance Early Career Fellowship [grant IA/E/21/1/506298], IIT Kanpur initiation grant [IITK /CS /2019236], Science and Engineering Research Board (SERB), Government of India Startup Research Grant [SRG/2020/001333], and Har Gobind Khorana Innovative Young Biotechnologist Award Grant [BT/13/IYBA/2020/05] to H.Z. A.S. Laboratory is supported by funding from NHMRC Ideas grant [2021/GNT2010795], MRFF EMCR grant [2022/MRF2016215], Perkins-Curtin start-up fellowship and Cancer Research Trust (CRT) Programme Grant to Liver Cancer Collaborative.

## Author Contributions

HZ and AS designed the study. HZ and NR developed the model and algorithm. NR implemented the software and performed all experiments and benchmarking. TK implemented the simulation experiment and performed benchmarking. RP, AM, FG and JC generated the fetal liver dataset. HZ, AS, and NR wrote the manuscript and all authors approved the manuscript.

## Competing Interests

The authors declare no competing interests.

## Notes

### Competing Interest Statement

The authors have declared no competing interest.

### Summary of Updates

The figures, main manuscript and supplementary material

