## Supplementary Material for "Charting spatial ligand-target activity using Renoir"

[1]Department of Computer Science and Engineering, Indian Institute of Technology Kanpur, India
[2]Department of Mathematics and Statistics, Indian Institute of Technology Kanpur, India
[3]Harry Perkins Institute of Medical Research, Nedlands, Perth, Western Australia
[4]Curtin Medical School, Curtin University, Perth, Western Australia
[5]University of Sydney, Sydney, Australia
[6]KK Research Center, KK Women's and Children's Hospital, Singapore
[7]Singapore Immunology Network (SIgN), Agency for Science, Technology and Research (ASTAR), Singapore
[8]Gustave Roussy Cancer Campus, Villejuif, France
[9]Translational Genomics Program, Garvan Institute of Medical Research, Darlinghurst, Australia
[10]School of Clinical Medicine, Faculty of Medicine and Health, University of New South Wales, Kensington, Australia
[11]Department of Biological Sciences and Bioengineering, Indian Institute of Technology Kanpur, India
[12]Mehta Family Center for Engineering in Medicine, Indian Institute of Technology Kanpur, India
[*]*Corresponding Author*
[3,4,6,9,10] (A.S.), [1,11,12] (H.Z.)

### Supplementary Figures

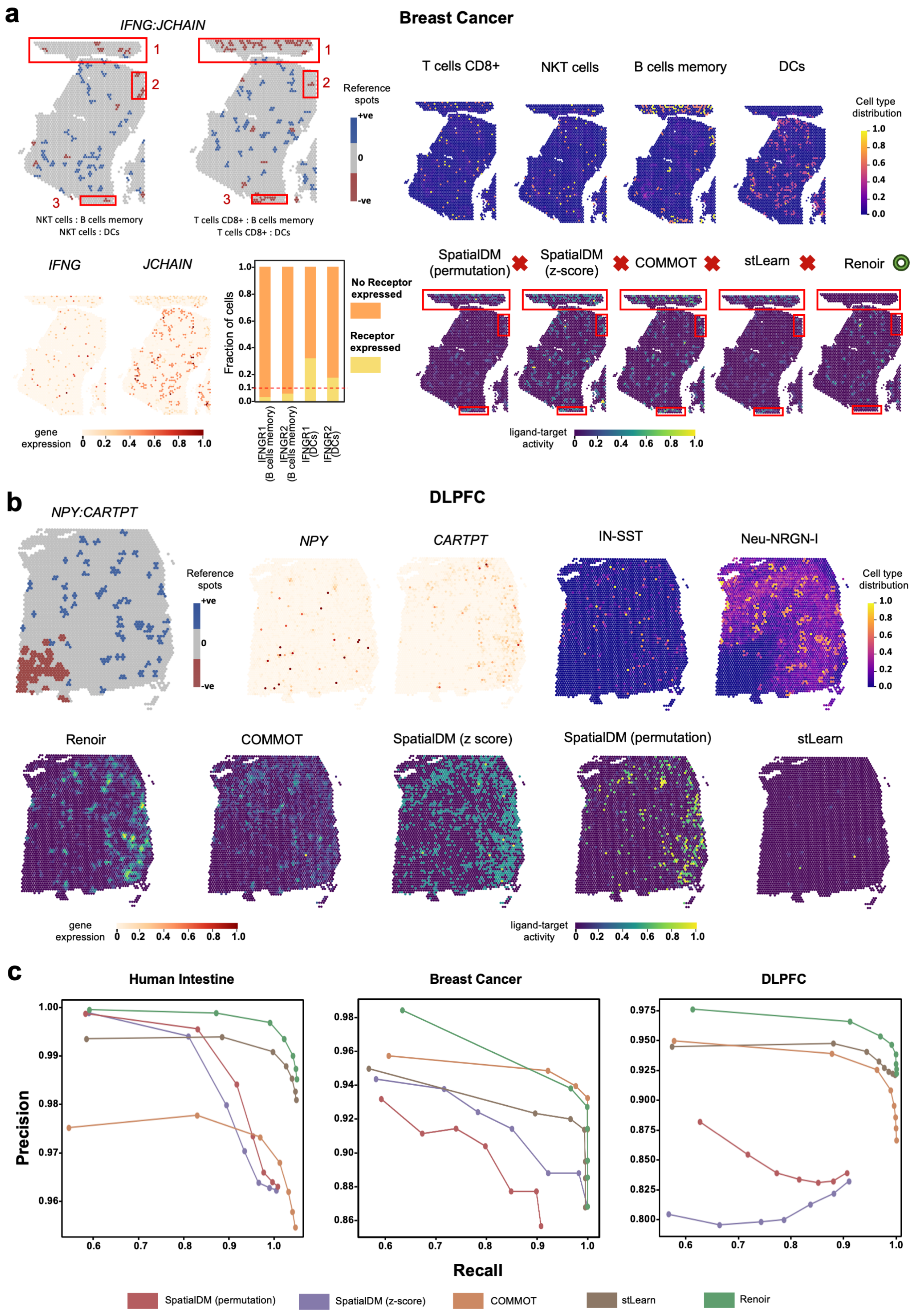

Supplementary Figure 1: (a) Comparison of Renoir, SpatialDM, COMMOT, and stLearn for the simulated dataset for the ligand-target pair *IFNG-JCHAIN* simulated based on the TNBC dataset. Regions 1, 2 and 3 contain ligand and target expression, however, these regions harbor B memory cells which do not express the receptor for *IFNG* (*IFNGR1* and *IFNGR2*) making it unfeasible for these regions to harbor *IFNG-JCHAIN* activity. (b) Comparison of Renoir, SpatialDM, COMMOT, and stLearn for the simulated dataset for the ligand-target pair *NPY-CARTPT* simulated based on the DLPFC dataset (c) Comparison of Renoir, SpatialDM, COMMOT, and stLearn in terms of average precision and recall over all the ligand-target pairs across the tissue types across different threshold values used for calculating precision and recall. For varying threshold values, Renoir was able to consistently maintain high precision and recall values across the tissue types.

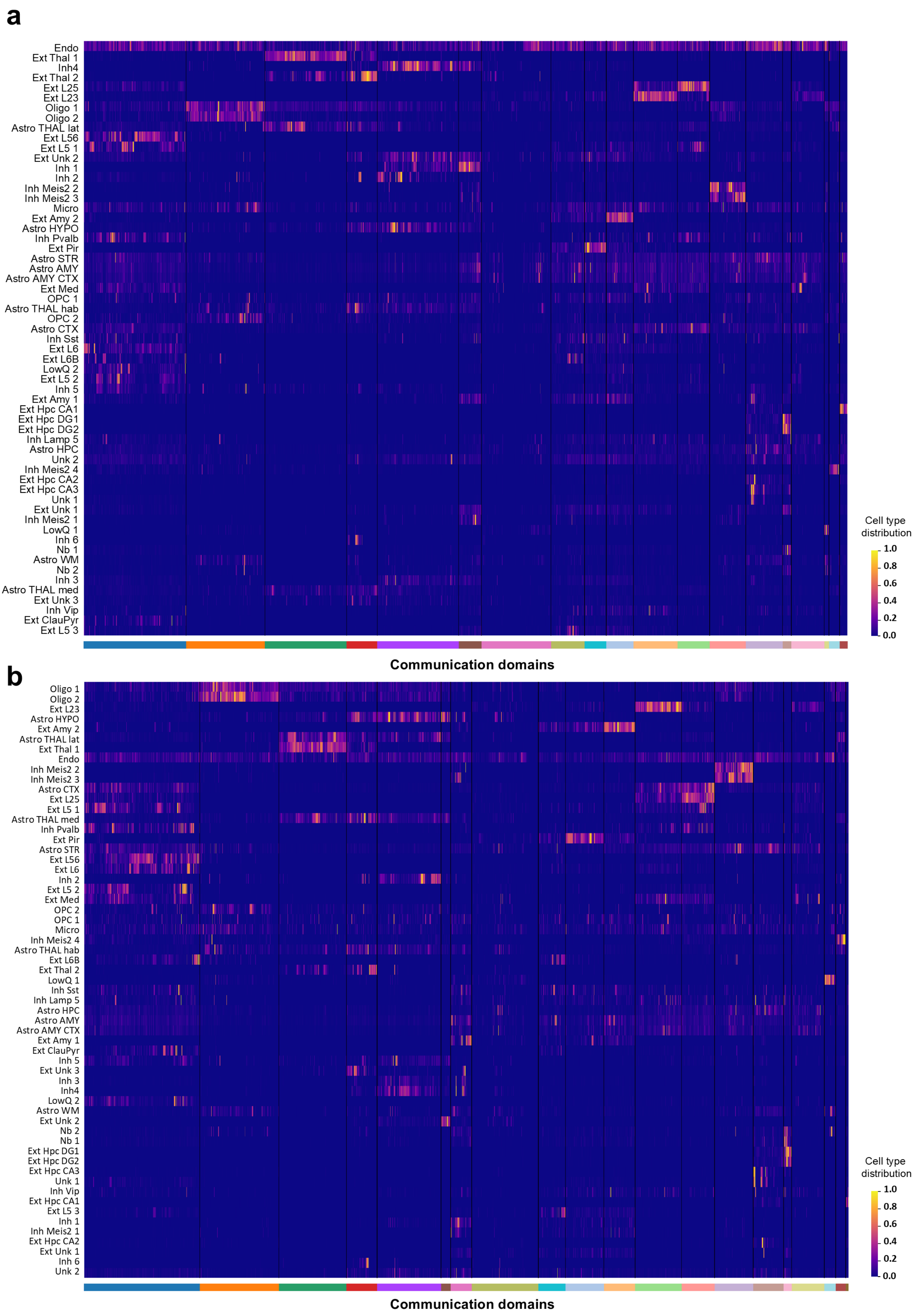


Supplementary Figure 2: Distribution of cell type abundances across the spots in mouse brain samples (a) S1 and (b) S2. The spots are ordered according to the spatial communication domains inferred by Renoir.

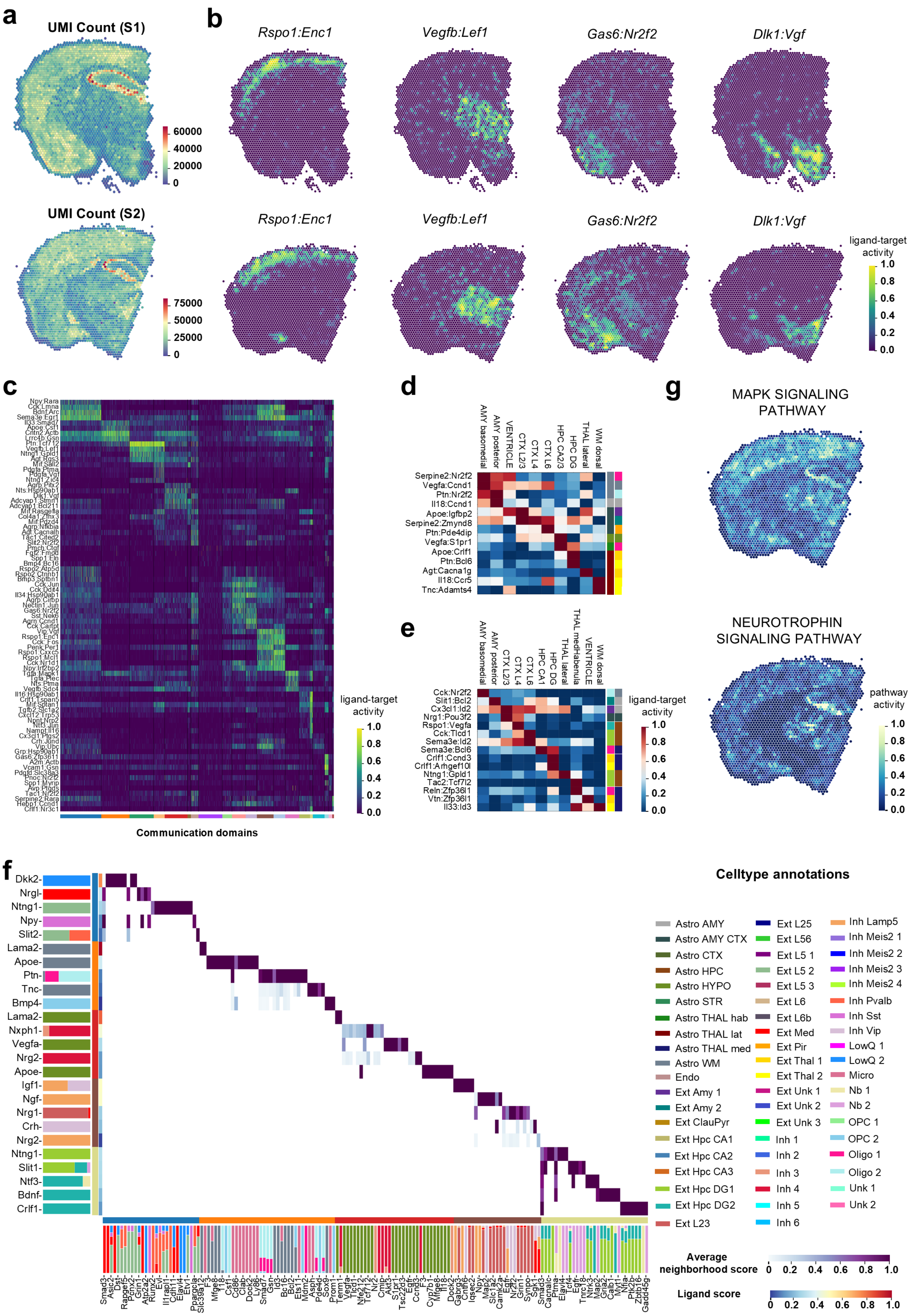
a
UMI Count (S1)
60000
40000
20000
0
UMI Count (S2)
75000
50000
25000
0
b
Rspo1:Enc1
Vegfb:Lef1
Gas6:Nr2f2
Dlk1:Vgf
Rspo1:Enc1
Vegfb:Lef1
Gas6:Nr2f2
Dlk1:Vgf
ligand-target activity
1.0
0.8
0.6
0.4
0.2
0.0
c
ligand-target activity
Communication domains
d
AMY basomedial
AMY posterior
VENTRICLE
CTX L2/3
CTX L4
CTX L6
HPC CA2/3
HPC DG
THAL lateral
WM dorsal
Serpine2:Nr2f2
Vegfa:Ccnd1
Ptn:Nr2f2
Il18:Ccnd1
Apoe:Igfbp2
Serpine2:Zmynd8
Ptn:Pde4dip
Vegfa:S1pr1
Apoe:Crlf1
Ptn:Bcl6
Agt:Cacna1g
Il18:Ccr5
Tnc:Adamts4
e
AMY basomedial
AMY posterior
CTX L2/3
CTX L4
CTX L6
HPC CA1
HPC DG
THAL lateral
THAL medhabenula
VENTRICLE
WM dorsal
Cck:Nr2f2
Slit1:Bcl2
Cx3cl1:Id2
Nrg1:Pou3f2
Rspo1:Vegfa
Cck:Tlcd1
Sema3e:Id2
Sema3e:Bcl6
Crlf1:Ccnd3
Crlf1:Arhgef10l
Ntng1:Gpld1
Tac2:Tcf7l2
Reln:Zfp36l1
Vtn:Zfp36l1
Il33:Id3
ligand-target activity
g
MAPK SIGNALING PATHWAY
NEUROTROPHIN SIGNALING PATHWAY
pathway activity
f
Dkk2-
Nrgl-
Ntng1-
Npy-
Slit2-
Lama2-
Apoe-
Ptn-
Tnc-
Bmp4-
Lama2-
Nxph1-
Vegfa-
Nrg2-
Apoe-
Igf1-
Ngf-
Nrg1-
Crh-
Nrg2-
Ntng1-
Slit1-
Ntf3-
Bdnf-
Crlf1-
Celltype annotations
Astro AMY
Astro AMY CTX
Astro CTX
Astro HPC
Astro HYPO
Astro STR
Astro THAL hab
Astro THAL lat
Astro THAL med
Astro WM
Endo
Ext Amy 1
Ext Amy 2
Ext ClauPyr
Ext Hpc CA1
Ext Hpc CA2
Ext Hpc CA3
Ext Hpc DG1
Ext Hpc DG2
Ext L23
Ext L25
Ext L56
Ext L5 1
Ext L5 2
Ext L5 3
Ext L6
Ext L6b
Ext Med
Ext Pir
Ext Thal 1
Ext Thal 2
Ext Unk 1
Ext Unk 2
Ext Unk 3
Inh 1
Inh 2
Inh 3
Inh 4
Inh 5
Inh 6
Inh Lamp5
Inh Meis2 1
Inh Meis2 2
Inh Meis2 3
Inh Meis2 4
Inh Pvalb
Inh Sst
Inh Vip
LowQ 1
LowQ 2
Micro
Nb 1
Nb 2
OPC 1
OPC 2
Oligo 1
Oligo 2
Unk 1
Unk 2
Average neighborhood score
0 0.2 0.4 0.6 0.8 1.0
Ligand score
0 0.2 0.4 0.6 0.8 1.0

Supplementary Figure 3: (a) Distribution of UMI count across mouse brain samples S1 and S2. (b) Spatial map of neighborhood activity scores for the ligand-target pairs across samples S1 and S2. (c) Differentially active ligand-target pairs inferred by Renoir across all communication domains for sample S2. (d) - (e) Characterization of ligand-target interactions between regional astrocyte subtypes and other co-localized cell types in sample S2. (d) Ligands expressed by regional astrocyte subtypes activating target genes in other cell types. (e) Target genes in regional astrocyte subtypes activated by ligands expressed by other co-localized cell types. (f) Communication domain-specific ranking of ligands based on cumulative activities over target genes expressed by major cell types in the domain for mouse brain sample S2. Top three ligands for each of eight communication domains are represented. Stacked color bars represent the cell types that express the ligand (right) and target (top). (g) Spatial map of MAPK SIGNALING and NEUROTROPHIN SIGNALING pathway activity across the spots in sample S2.

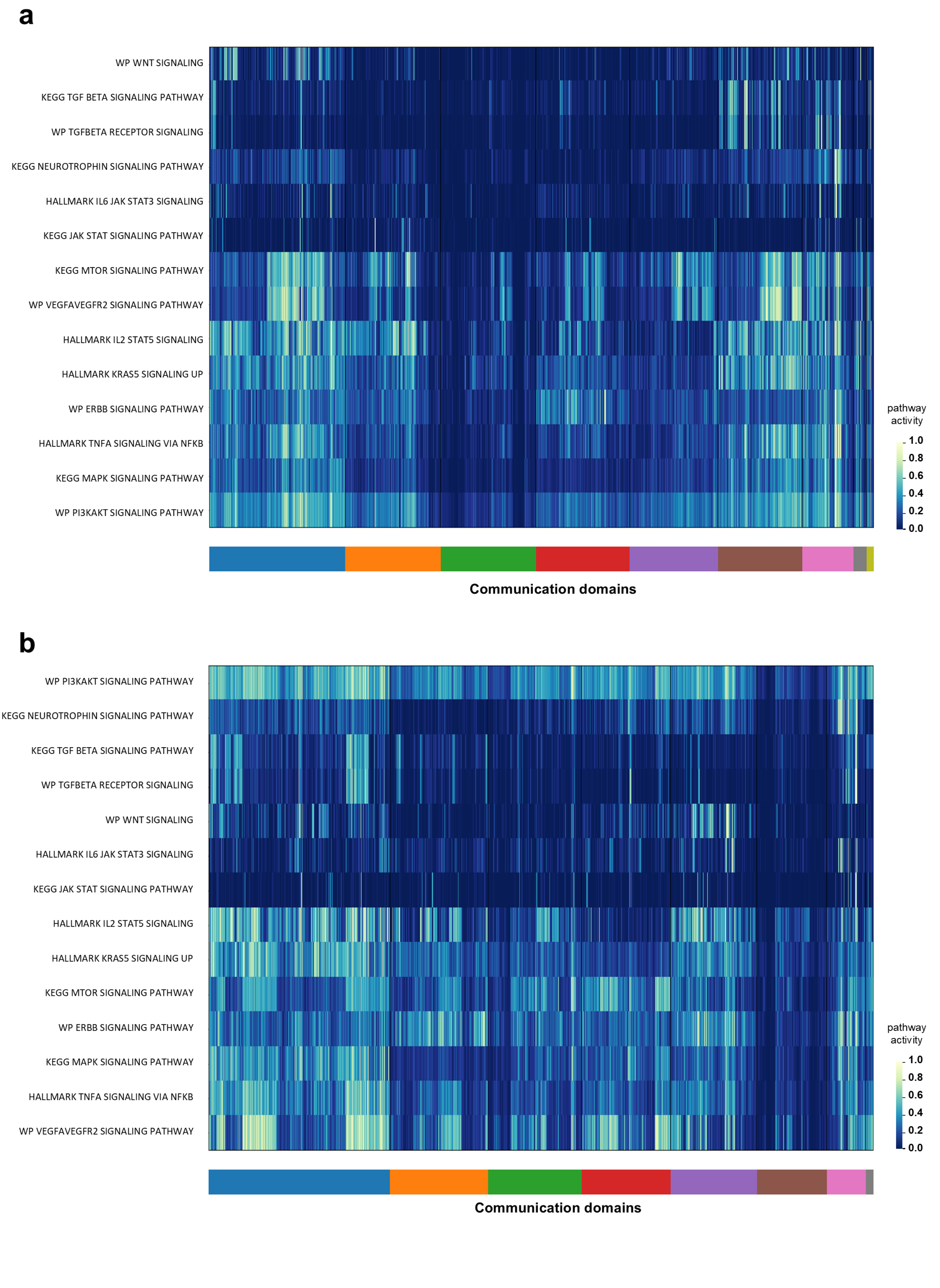


Supplementary Figure 4: (a) - (b) Distribution of various pathway activity across the spots in mouse brain samples (a) S1 and (b) S2. The spots are ordered according to the spatial communication domains inferred by Renoir.

a

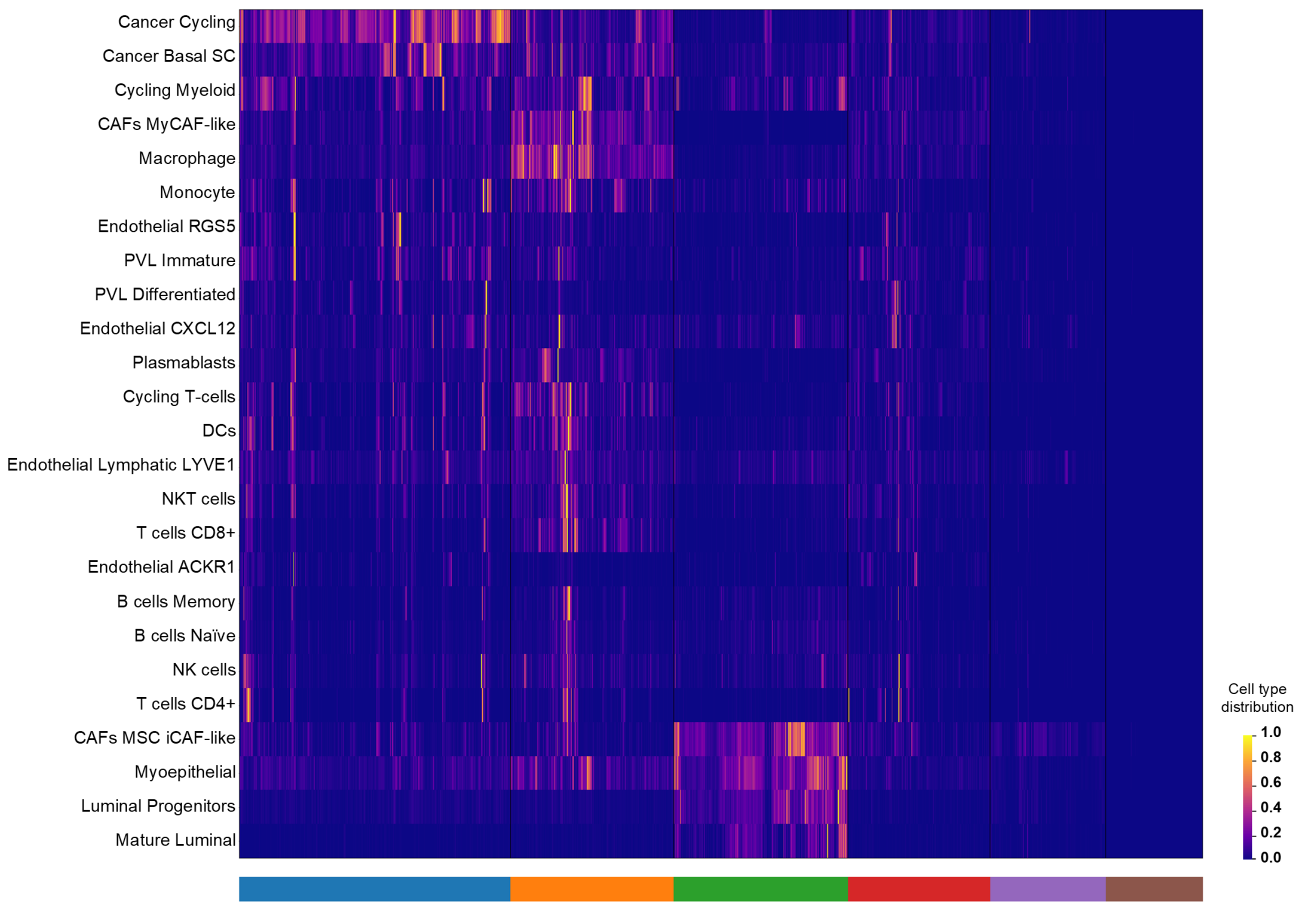


b

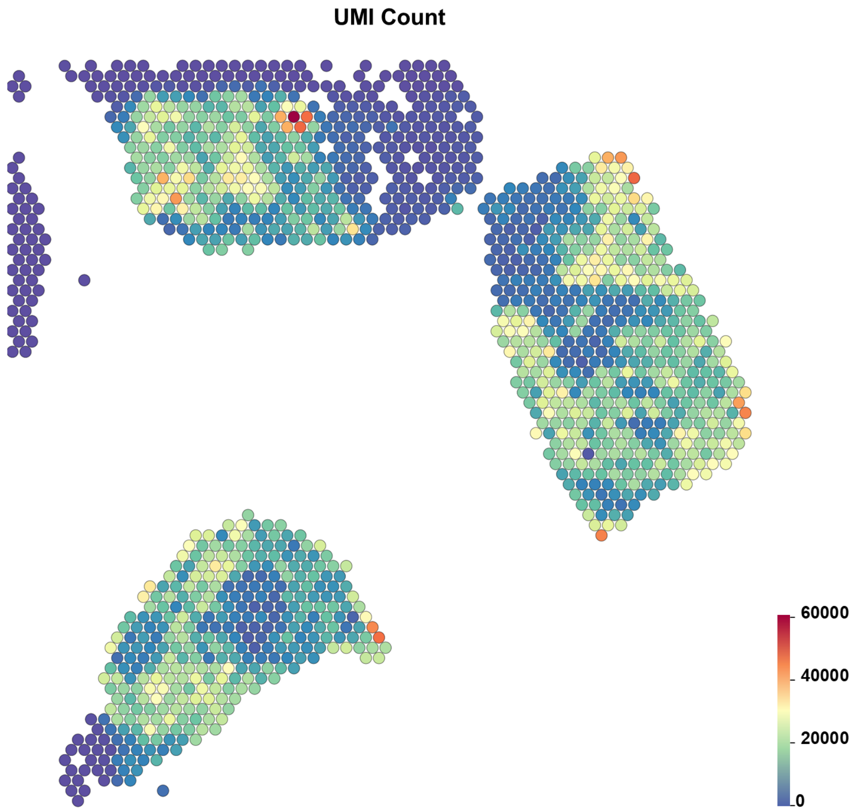


Supplementary Figure 5: (a) Distribution of cell types across spots in TNBC sample (b) Distribution of UMI count across the TNBC sample.

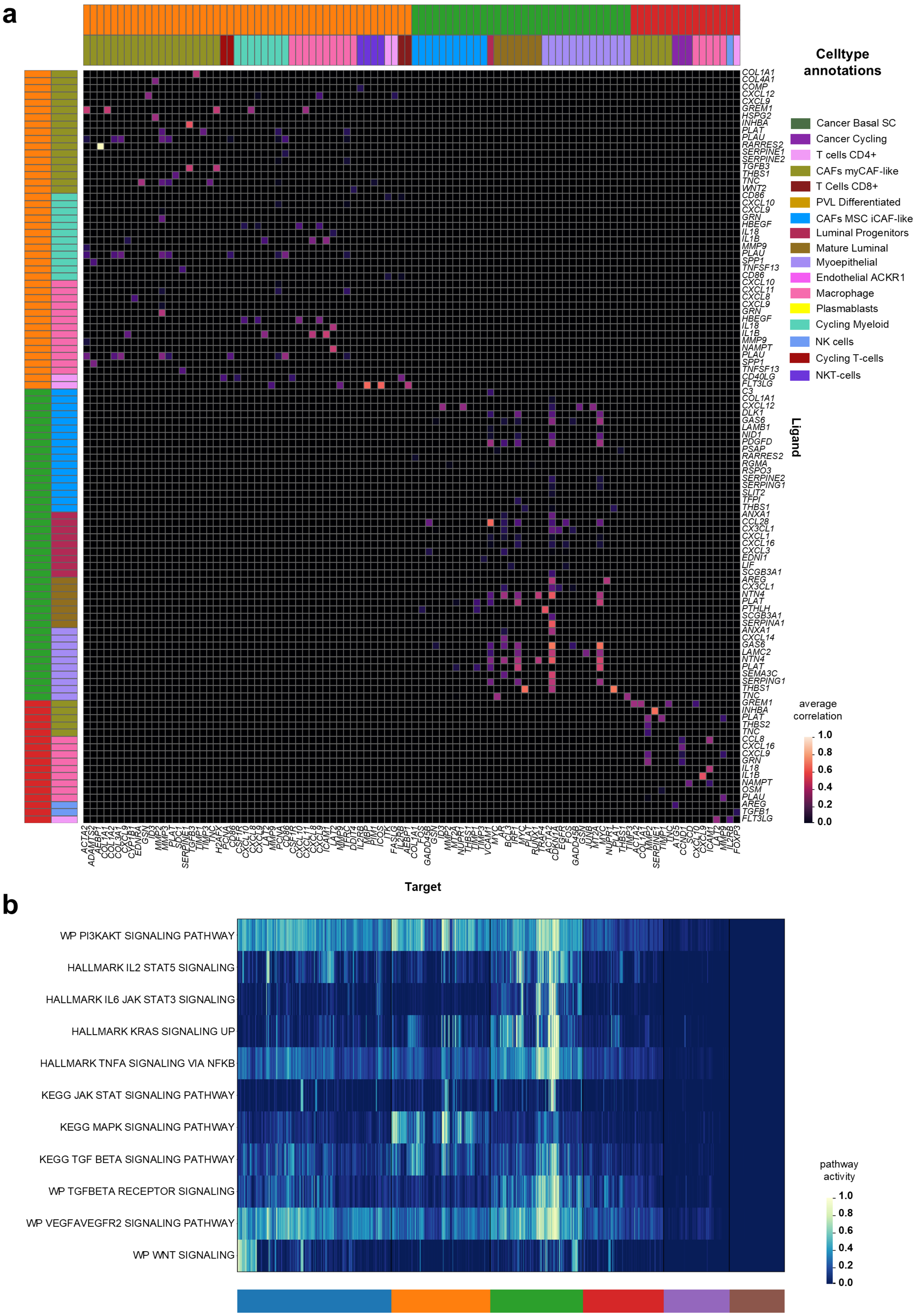
a
Celltype annotations
Cancer Basal SC
Cancer Cycling
T cells CD4+
CAFs myCAF-like
T Cells CD8+
PVL Differentiated
CAFs MSC iCAF-like
Luminal Progenitors
Mature Luminal
Myoepithelial
Endothelial ACKR1
Macrophage
Plasmablasts
Cycling Myeloid
NK cells
Cycling T-cells
NKT-cells
Ligand
average correlation
1.0
0.8
0.6
0.4
0.2
0.0
Target
b
WP PI3KAKT SIGNALING PATHWAY
HALLMARK IL2 STAT5 SIGNALING
HALLMARK IL6 JAK STAT3 SIGNALING
HALLMARK KRAS SIGNALING UP
HALLMARK TNFA SIGNALING VIA NFKB
KEGG JAK STAT SIGNALING PATHWAY
KEGG MAPK SIGNALING PATHWAY
KEGG TGF BETA SIGNALING PATHWAY
WP TGFBETA RECEPTOR SIGNALING
WP VEGFAVEGFR2 SIGNALING PATHWAY
WP WNT SIGNALING
pathway activity
1.0
0.8
0.6
0.4
0.2
0.0

Supplementary Figure 6: (a) Cell type-specific ligand-target interactions inferred by Renoir for the TNBC dataset for the major cell types in domains 1, 2, and 3. The inner color bar represents the cell types of the ligand (left) and target (top) and the outer color bar represents the communication domain. Each cell represents the average Pearson correlation between the ligand-target neighborhood scores and the abundances of the cell types expressing the ligand and target across the spots pertaining to the domain being considered. (b) Distribution of pathway activities across the communication domains in the TNBC sample.

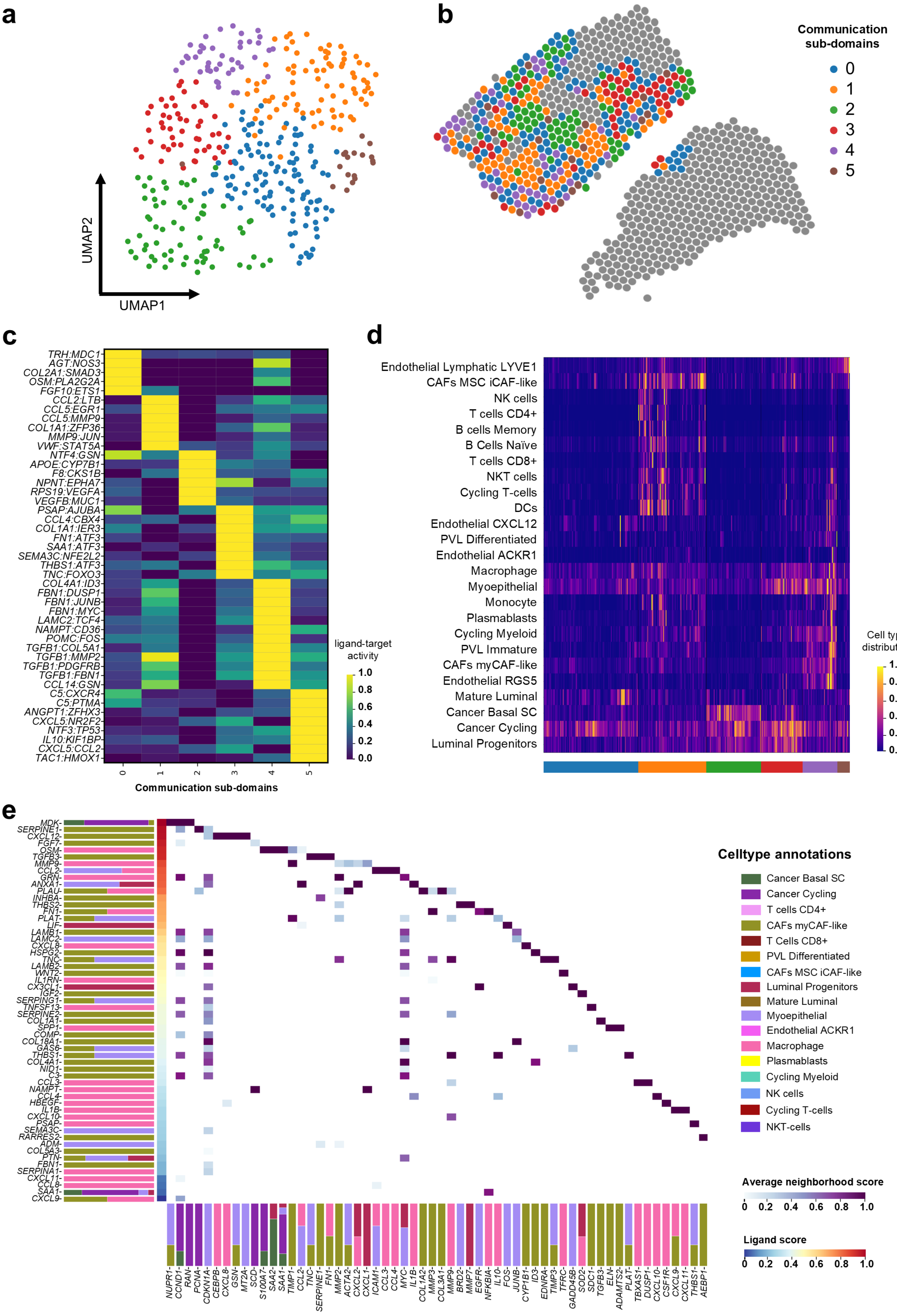

a
UMAP2
UMAP1
b
Communication sub-domains
0
1
2
3
4
5
c
TRH:MDC1
AGT:NOS3
COL2A1:SMAD3
OSM:PLA2G2A
FGF10:ETS1
CCL2:LTB
CCL5:EGR1
CCL5:MMP9
COL1A1:ZFP36
MMP9:JUN
VWF:STAT5A
NTF4:GSN
APOE:CYP7B1
F8:CKS1B
NPNT:EPHA7
RPS19:VEGFA
VEGFB:MUC1
PSAP:AJUBA
CCL4:CBX4
COL1A1:IER3
FN1:ATF3
SAA1:ATF3
SEMA3C:NFE2L2
THBS1:ATF3
TNC:FOXO3
COL4A1:ID3
FBN1:DUSP1
FBN1:JUNB
FBN1:MYC
LAMC2:TCF4
NAMPT:CD36
POMC:FOS
TGFB1:COL5A1
TGFB1:MMP2
TGFB1:PDGFRB
TGFB1:FBN1
CCL14:GSN
C5:CXCR4
C5:PTMA
ANGPT1:ZFHX3
CXCL5:NR2F2
NTF3:TP53
IL10:KIF1BP
CXCL5:CCL2
TAC1:HMOX1
ligand-target activity
1.0
0.8
0.6
0.4
0.2
0.0
0
1
2
3
4
5
Communication sub-domains
d
Endothelial Lymphatic LYVE1
CAFs MSC iCAF-like
NK cells
T cells CD4+
B cells Memory
B Cells Naïve
T cells CD8+
NKT cells
Cycling T-cells
DCs
Endothelial CXCL12
PVL Differentiated
Endothelial ACKR1
Macrophage
Myoepithelial
Monocyte
Plasmablasts
Cycling Myeloid
PVL Immature
CAFs myCAF-like
Endothelial RGS5
Mature Luminal
Cancer Basal SC
Cancer Cycling
Luminal Progenitors
Cell type distribution
1.0
0.8
0.6
0.4
0.2
0.0
e
MDK
SERPINE1
CXCL12
FGF7
OSM
TGFB3
MMP9
CCL2
GRN
ANXA1
PLAU
INHBA
THBS2
FN1
PLAT
LIF
LAMB1
LAMC2
CXCL8
HSPG2
TNC
LAMB2
WNT2
IL1RN
CX3CL1
IGF2
SERPING1
TNFSF13
SERPINE2
COL1A1
SPP1
COMP
COL18A1
GAS6
THBS1
COL4A1
NID1
C3
CCL3
NAMPT
CCL4
HBEGF
IL1B
CXCL10
PSAP
SEMA3C
RARRES2
ADM
COL5A3
PTN
FBN1
SERPINA1
CXCL11
CCL8
SAA1
CXCL9
NUPR1
CCND1
RAN
PCNA
CDKN1A
CEBPB
CXCL8
GSN
MT2A
SCD
S100A7
SAA2
SAA1
TIMP1
CCL2
TNC
SERPINE1
FN1
MMP2
ACTA2
CXCL2
CXCL1
ICAM1
CCL3
CCL4
MYC
IL1B
COL1A2
MMP3
COL3A1
MMP9
BRD2
MMP7
EGFR
NFKBIA
IL10
FOS
JUNB
CYP1B1
ID3
EDNRA
TIMP3
TFRC
GADD45B
SOD2
SDC1
TGFB3
ELN
ADAMTS2
PLAT
TBXAS1
DUSP1
CXCL10
CSF1R
CXCL9
CXCL11
THBS1
AEBP1
Celltype annotations
Cancer Basal SC
Cancer Cycling
T cells CD4+
CAFs myCAF-like
T Cells CD8+
PVL Differentiated
CAFs MSC iCAF-like
Luminal Progenitors
Mature Luminal
Myoepithelial
Endothelial ACKR1
Macrophage
Plasmablasts
Cycling Myeloid
NK cells
Cycling T-cells
NKT-cells
Average neighborhood score
0
0.2
0.4
0.6
0.8
1.0
Ligand score
0
0.2
0.4
0.6
0.8
1.0

Supplementary Figure 7: (a) UMAP embeddings and (b) spatial map of communication sub-domains of domain 0 of the TNBC sample. (c) Differentially active ligand-target pairs across the communication sub-domains of domain 0. (d) Inferred cell type distribution across the communication sub-domains. (e) Ranking of ligands based on their cumulative activities over target genes expressed by major cell types in domain 0.

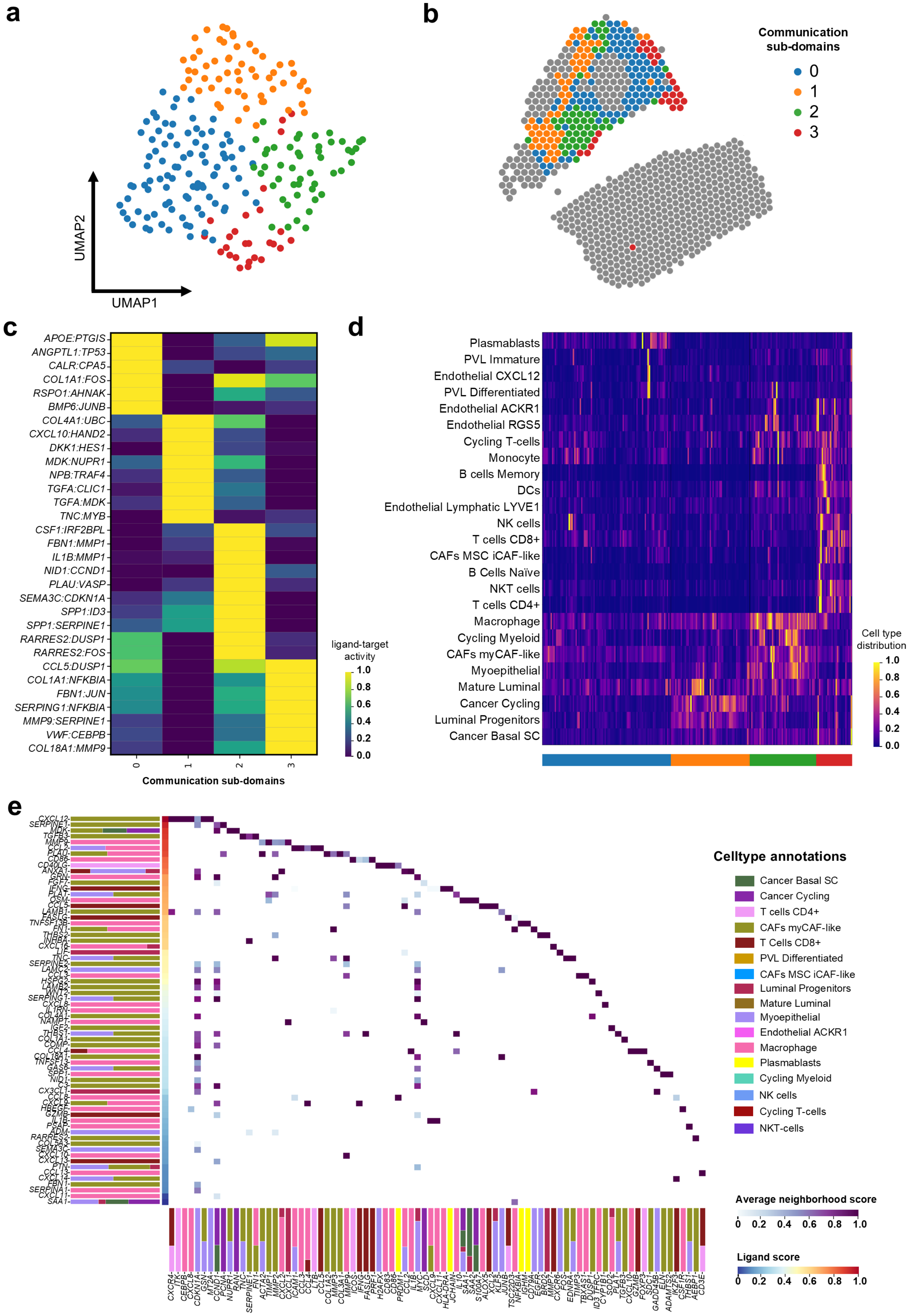
a
UMAP2
UMAP1
b
Communication sub-domains
0
1
2
3
c
APOE:PTGIS
ANGPTL1:TP53
CALR:CPA5
COL1A1:FOS
RSPO1:AHNAK
BMP6:JUNB
COL4A1:UBC
CXCL10:HAND2
DKK1:HES1
MDK:NUPR1
NPB:TRAF4
TGFA:CLIC1
TGFA:MDK
TNC:MYB
CSF1:IRF2BPL
FBN1:MMP1
IL1B:MMP1
NID1:CCND1
PLAU:VASP
SEMA3C:CDKN1A
SPP1:ID3
SPP1:SERPINE1
RARRES2:DUSP1
RARRES2:FOS
CCL5:DUSP1
COL1A1:NFKBIA
FBN1:JUN
SERPING1:NFKBIA
MMP9:SERPINE1
VWF:CEBPB
COL18A1:MMP9
0
1
2
3
Communication sub-domains
ligand-target activity
1.0
0.8
0.6
0.4
0.2
0.0
d
Plasmablasts
PVL Immature
Endothelial CXCL12
PVL Differentiated
Endothelial ACKR1
Endothelial RGS5
Cycling T-cells
Monocyte
B cells Memory
DCs
Endothelial Lymphatic LYVE1
NK cells
T cells CD8+
CAFs MSC iCAF-like
B Cells Naïve
NKT cells
T cells CD4+
Macrophage
Cycling Myeloid
CAFs myCAF-like
Myoepithelial
Mature Luminal
Cancer Cycling
Luminal Progenitors
Cancer Basal SC
Cell type distribution
1.0
0.8
0.6
0.4
0.2
0.0
e
Celltype annotations
Cancer Basal SC
Cancer Cycling
T cells CD4+
CAFs myCAF-like
T Cells CD8+
PVL Differentiated
CAFs MSC iCAF-like
Luminal Progenitors
Mature Luminal
Myoepithelial
Endothelial ACKR1
Macrophage
Plasmablasts
Cycling Myeloid
NK cells
Cycling T-cells
NKT-cells
Average neighborhood score
0 0.2 0.4 0.6 0.8 1.0
Ligand score
0 0.2 0.4 0.6 0.8 1.0

Supplementary Figure 8: (a) UMAP embeddings and (b) spatial map of communication sub-domains of domain 1 of the TNBC sample. (c) Differentially active ligand-target pairs across the communication sub-domains of domain 1. (d) Inferred cell type distribution across the communication sub-domains. (e) Ranking of ligands based on their cumulative activities over target genes expressed by major cell types in domain 1.

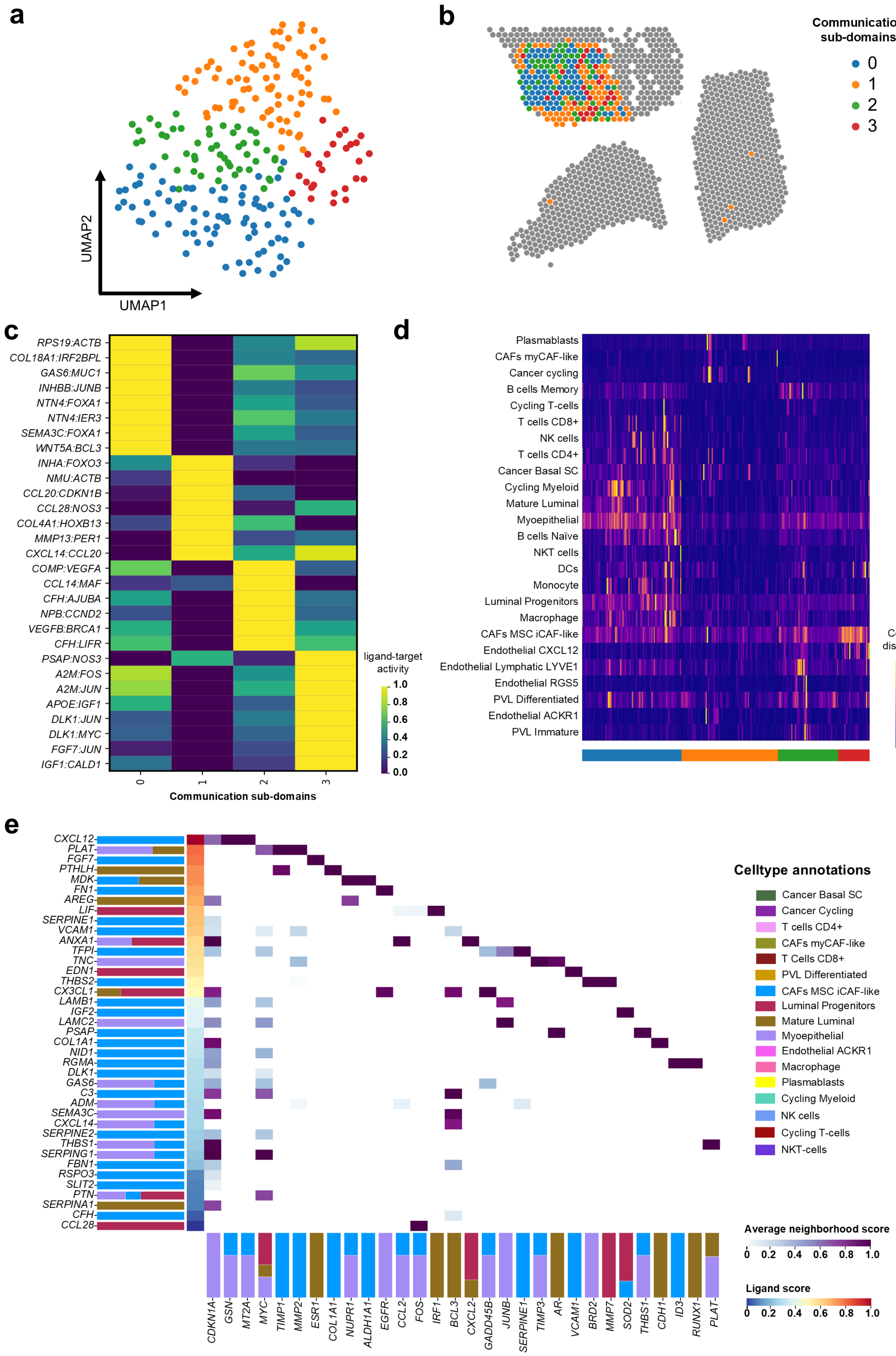

a
UMAP1
UMAP2
b
Communication sub-domains
0
1
2
3
c
RPS19:ACTB
COL18A1:IRF2BPL
GAS6:MUC1
INHBB:JUNB
NTN4:FOXA1
NTN4:IER3
SEMA3C:FOXA1
WNT5A:BCL3
INHA:FOXO3
NMU:ACTB
CCL20:CDKN1B
CCL28:NOS3
COL4A1:HOXB13
MMP13:PER1
CXCL14:CCL20
COMP:VEGFA
CCL14:MAF
CFH:AJUBA
NPB:CCND2
VEGFB:BRCA1
CFH:LIFR
PSAP:NOS3
A2M:FOS
A2M:JUN
APOE:IGF1
DLK1:JUN
DLK1:MYC
FGF7:JUN
IGF1:CALD1
Communication sub-domains
ligand-target activity
1.0
0.8
0.6
0.4
0.2
0.0
d
Plasmablasts
CAFs myCAF-like
Cancer cycling
B cells Memory
Cycling T-cells
T cells CD8+
NK cells
T cells CD4+
Cancer Basal SC
Cycling Myeloid
Mature Luminal
Myoepithelial
B cells Naïve
NKT cells
DCs
Monocyte
Luminal Progenitors
Macrophage
CAFs MSC iCAF-like
Endothelial CXCL12
Endothelial Lymphatic LYVE1
Endothelial RGS5
PVL Differentiated
Endothelial ACKR1
PVL Immature
Cell type distribution
e
CXCL12
PLAT
FGF7
PTHLH
MDK
FN1
AREG
LIF
SERPINE1
VCAM1
ANXA1
TFPI
TNC
EDN1
THBS2
CX3CL1
LAMB1
IGF2
LAMC2
PSAP
COL1A1
NID1
RGMA
DLK1
GAS6
C3
ADM
SEMA3C
CXCL14
SERPINE2
THBS1
SERPING1
FBN1
RSPO3
SLIT2
PTN
SERPINA1
CFH
CCL28
CDKN1A
GSN
MT2A
MYC
TIMP1
MMP2
ESR1
COL1A1
NUPR1
ALDH1A1
EGFR
CCL2
FOS
IRF1
BCL3
CXCL2
GADD45B
JUNB
SERPINE1
TIMP3
AR
VCAM1
BRD2
MMP7
SOD2
THBS1
CDH1
ID3
RUNX1
PLAT
Celltype annotations
Cancer Basal SC
Cancer Cycling
T cells CD4+
CAFs myCAF-like
T Cells CD8+
PVL Differentiated
CAFs MSC iCAF-like
Luminal Progenitors
Mature Luminal
Myoepithelial
Endothelial ACKR1
Macrophage
Plasmablasts
Cycling Myeloid
NK cells
Cycling T-cells
NKT-cells
Average neighborhood score
Ligand score
0 0.2 0.4 0.6 0.8 1.0

Supplementary Figure 9: (a) UMAP embeddings and (b) spatial map of communication sub-domains of domain 2 of the TNBC sample. (c) Differentially active ligand-target pairs across the communication sub-domains of domain 2. (d) Inferred cell type distribution across the communication sub-domains. (e) Ranking of ligands based on their cumulative activities over target genes expressed by major cell types in domain 2.

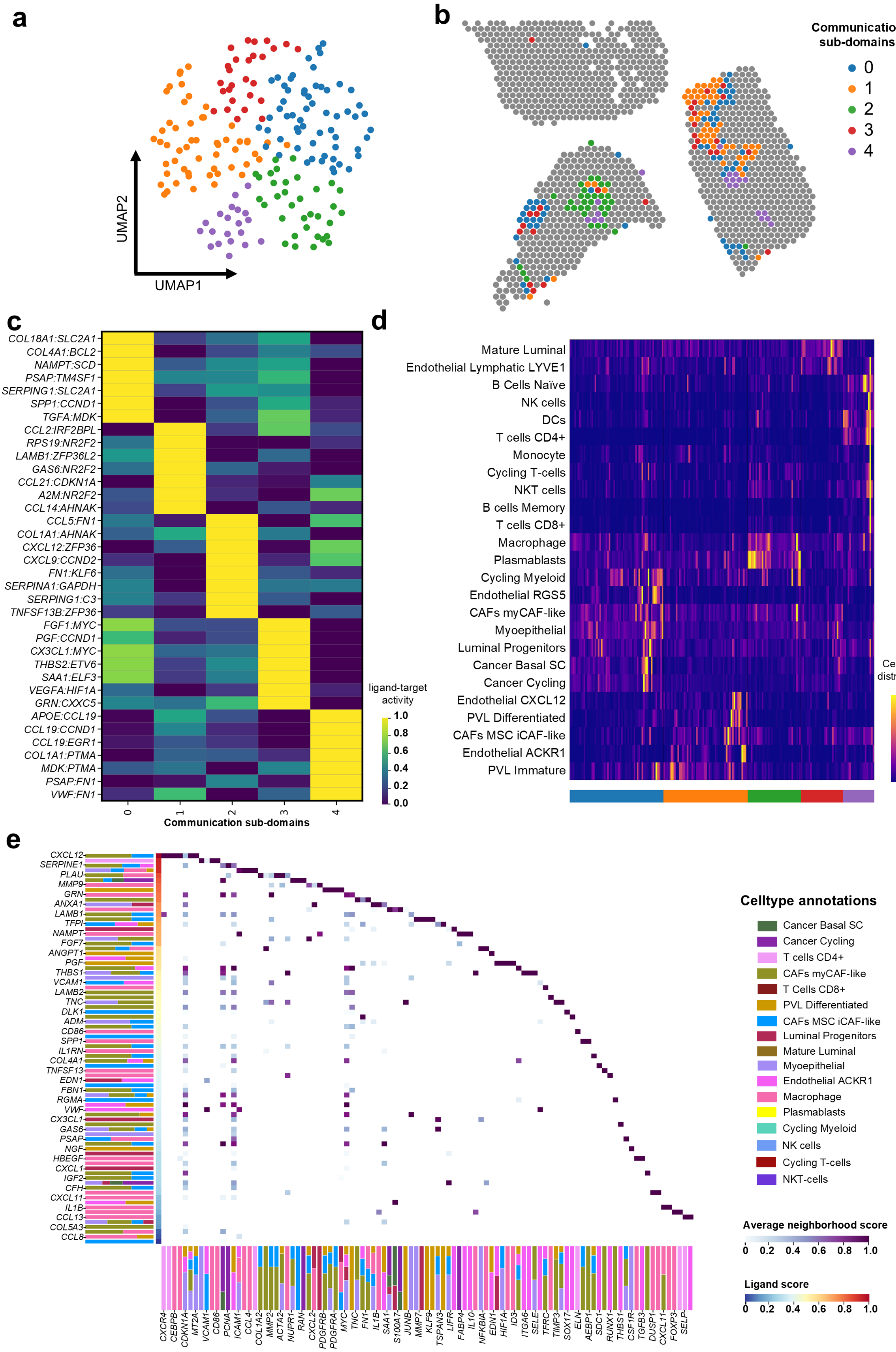

a
UMAP2
UMAP1
b
Communication sub-domains
0
1
2
3
4
c
COL18A1:SLC2A1
COL4A1:BCL2
NAMPT:SCD
PSAP:TM4SF1
SERPING1:SLC2A1
SPP1:CCND1
TGFA:MDK
CCL2:IRF2BPL
RPS19:NR2F2
LAMB1:ZFP36L2
GAS6:NR2F2
CCL21:CDKN1A
A2M:NR2F2
CCL14:AHNAK
CCL5:FN1
COL1A1:AHNAK
CXCL12:ZFP36
CXCL9:CCND2
FN1:KLF6
SERPINA1:GAPDH
SERPING1:C3
TNFSF13B:ZFP36
FGF1:MYC
PGF:CCND1
CX3CL1:MYC
THBS2:ETV6
SAA1:ELF3
VEGFA:HIF1A
GRN:CXXC5
APOE:CCL19
CCL19:CCND1
CCL19:EGR1
COL1A1:PTMA
MDK:PTMA
PSAP:FN1
VWF:FN1
0
1
2
3
4
Communication sub-domains
ligand-target activity
1.0
0.8
0.6
0.4
0.2
0.0
d
Mature Luminal
Endothelial Lymphatic LYVE1
B Cells Naïve
NK cells
DCs
T cells CD4+
Monocyte
Cycling T-cells
NKT cells
B cells Memory
T cells CD8+
Macrophage
Plasmablasts
Cycling Myeloid
Endothelial RGS5
CAFs myCAF-like
Myoepithelial
Luminal Progenitors
Cancer Basal SC
Cancer Cycling
Endothelial CXCL12
PVL Differentiated
CAFs MSC iCAF-like
Endothelial ACKR1
PVL Immature
Cell type distribution
1.0
0.8
0.6
0.4
0.2
0.0
e
CXCL12
SERPINE1
PLAU
MMP9
GRN
ANXA1
LAMB1
TFPI
NAMPT
FGF7
ANGPT1
PGF
THBS1
VCAM1
LAMB2
TNC
DLK1
ADM
CD86
SPP1
IL1RN
COL4A1
TNFSF13
EDN1
FBN1
RGMA
VWF
CX3CL1
GAS6
PSAP
NGF
HBEGF
CXCL1
IGF2
CFH
CXCL11
IL1B
CCL13
COL5A3
CCL8
CXCR4
CEBPB
CDKN1A
MT2A
VCAM1
CD86
PCNA
ICAM1
CCL4
COL1A2
MMP2
ACTA2
NUPR1
RAN
CXCL2
PDGFRB
PDGFRA
MYC
TNC
FN1
IL1B
SAA1
S100A7
JUNB
MMP7
KLF9
TSPAN3
LIFR
FABP4
IL10
NFKBIA
EDN1
HIF1A
ID3
ITGA6
SELE
TFRC
TIMP3
SOX17
ELN
AEBP1
SDC1
RUNX1
THBS1
CSF1R
TGFB3
DUSP1
CXCL11
FOXP3
SELP
Celltype annotations
Cancer Basal SC
Cancer Cycling
T cells CD4+
CAFs myCAF-like
T Cells CD8+
PVL Differentiated
CAFs MSC iCAF-like
Luminal Progenitors
Mature Luminal
Myoepithelial
Endothelial ACKR1
Macrophage
Plasmablasts
Cycling Myeloid
NK cells
Cycling T-cells
NKT-cells
Average neighborhood score
0
0.2
0.4
0.6
0.8
1.0
Ligand score
0
0.2
0.4
0.6
0.8
1.0

Supplementary Figure 10: (a) UMAP embeddings and (b) spatial map of communication sub-domains of domain 3 of the TNBC sample. (c) Differentially active ligand-target pairs across the communication sub-domains of domain 3. (d) Inferred cell type distribution across the communication sub-domains. (e) Ranking of ligands based on their cumulative activities over target genes expressed by major cell types in domain 3.

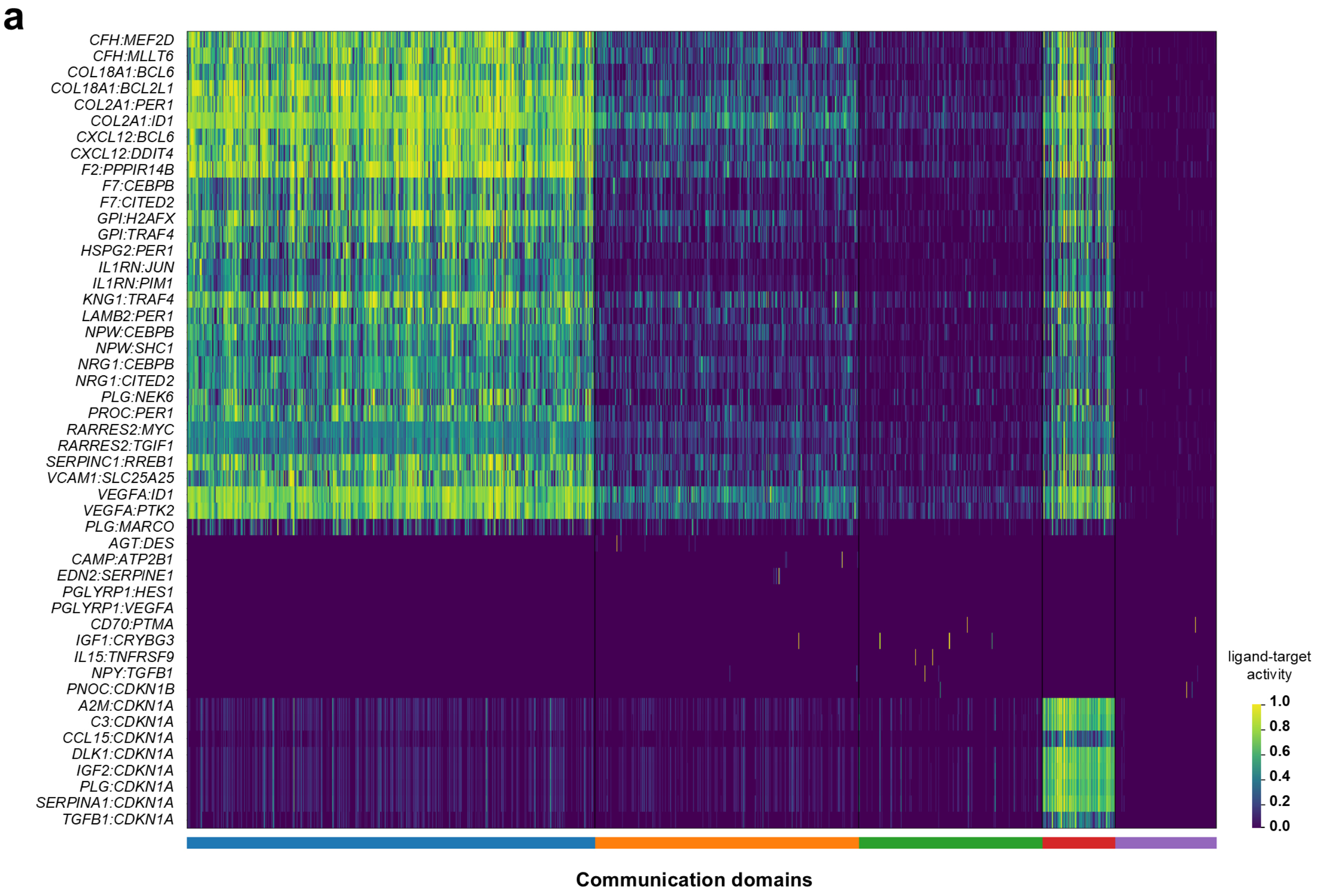


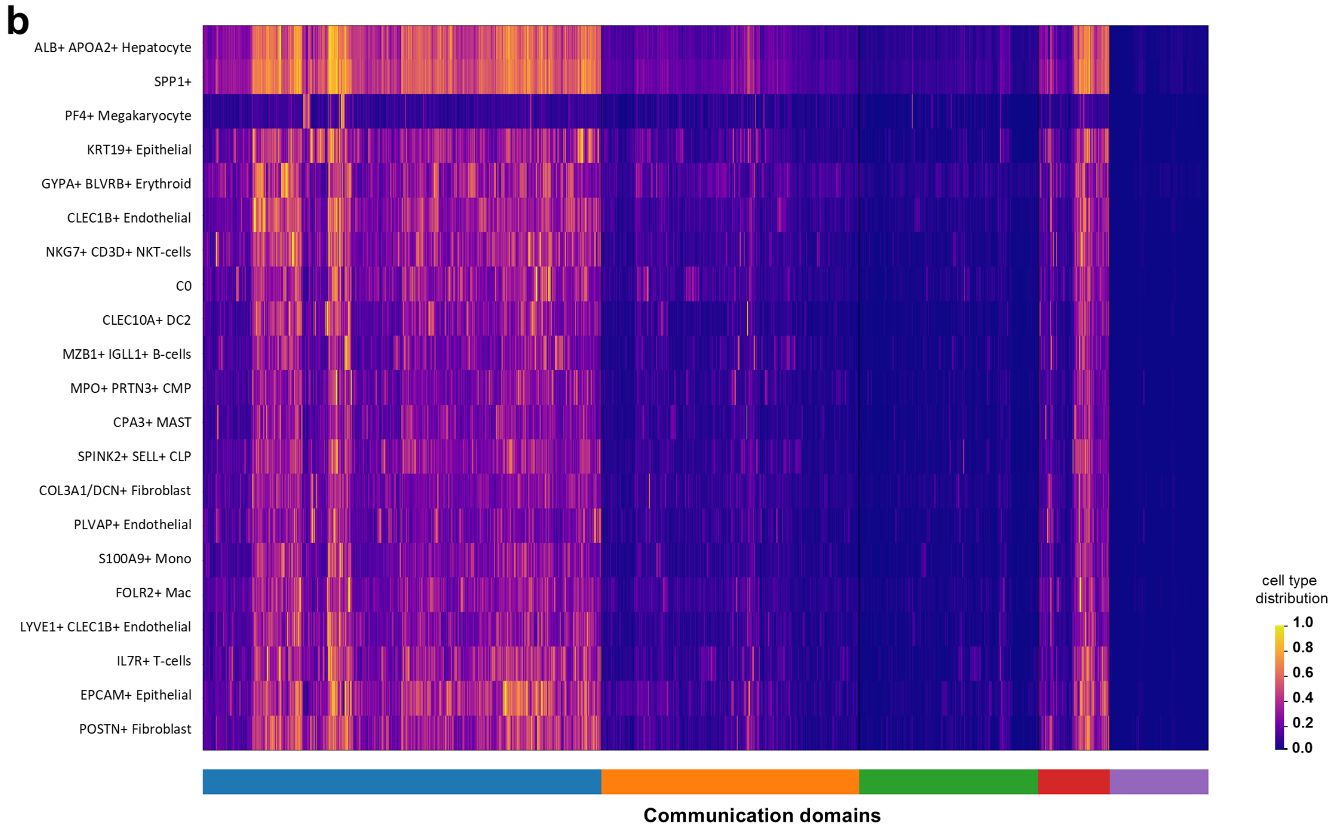


Supplementary Figure 11: (a) Differentially active ligand-target pairs inferred by Renoir across all communication domains for the fetal liver dataset. (b) Distribution of cell types across spots in the fetal liver dataset.

**a**

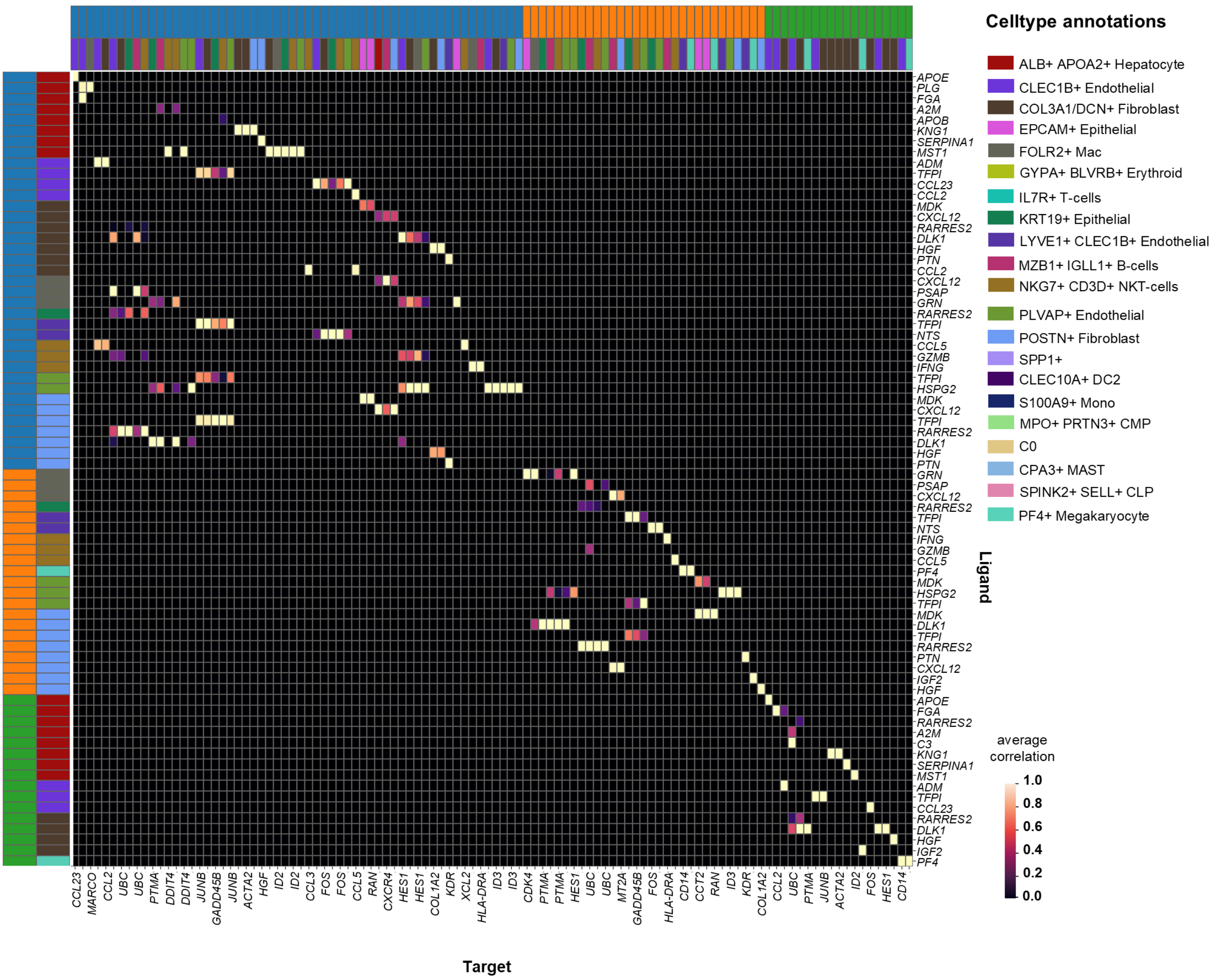

**b**

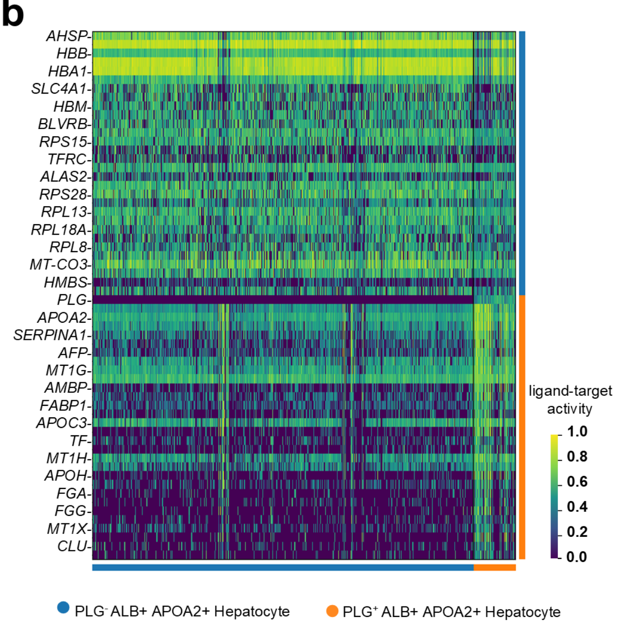
AHSP
HBB
HBA1
SLC4A1
HBM
BLVRB
RPS15
TFRC
ALAS2
RPS28
RPL13
RPL18A
RPL8
MT-CO3
HMBS
PLG
APOA2
SERPINA1
AFP
MT1G
AMBP
FABP1
APOC3
TF
MT1H
APOH
FGA
FGG
MT1X
CLU
ligand-target activity
1.0
0.8
0.6
0.4
0.2
0.0
PLG- ALB+ APOA2+ Hepatocyte
PLG+ ALB+ APOA2+ Hepatocyte

Supplementary Figure 12: (a) Cell type-specific ligand-target interactions inferred by Renoir for the major cell types in domains 0, 1, and 2. The inner color bar represents the cell types of the ligand (left) and target (top) and the outer color bar represents the communication domain. Each cell represents the average Pearson correlation between the ligand-target neighborhood scores and the abundances of the cell types expressing the ligand and target across the spots pertaining to the domain being considered. (b) Differentially expressed genes between $PLG^{+}$ and $PLG^{-}$ Hepatocyte populations.

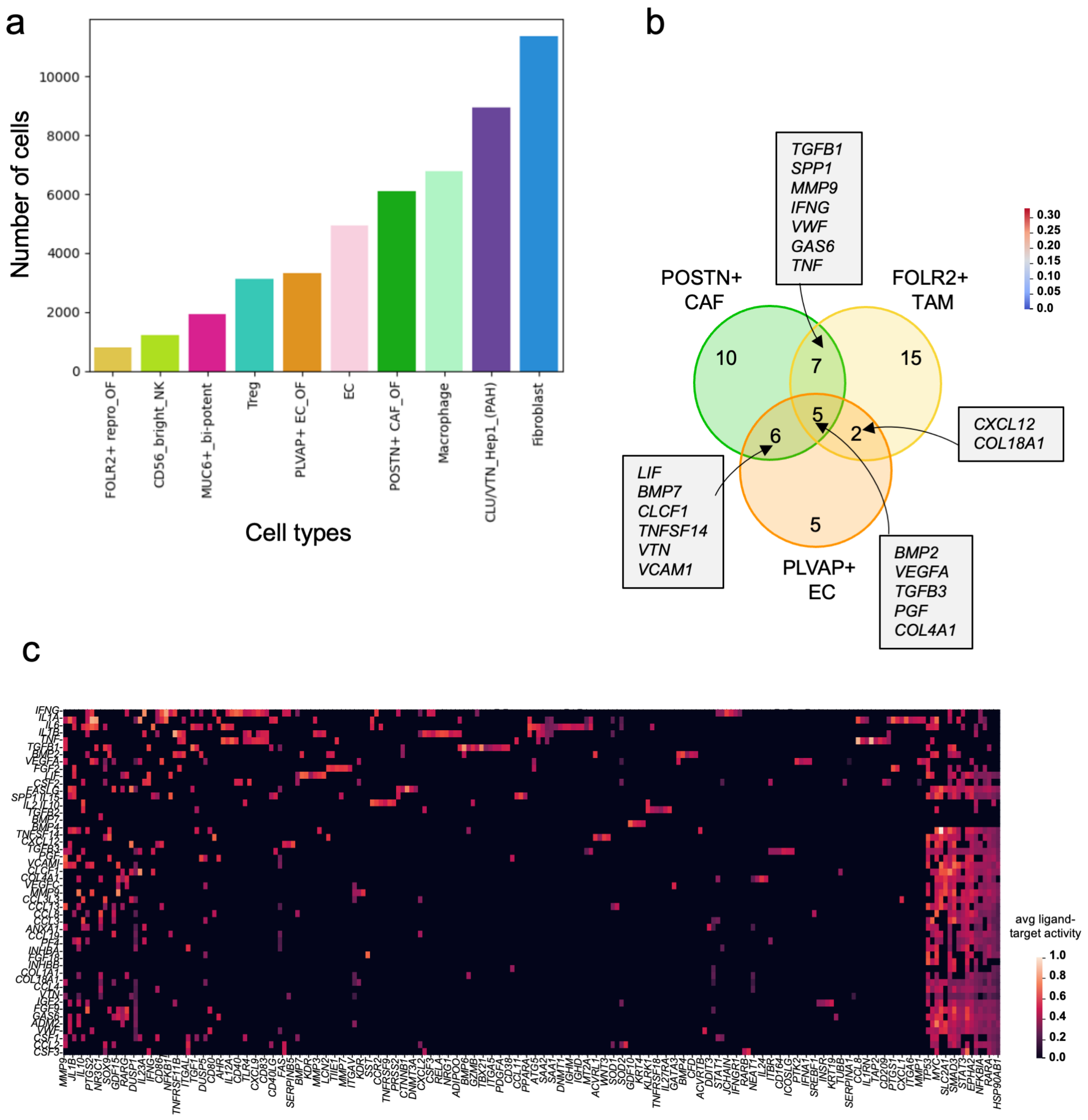


Supplementary Figure 13: (a) Distribution of major cell types located in the neighborhoods of onco-fetal cell types for the Nanostring CosMx hepatocellular carcinoma dataset. Cell types are ranked in the ascending order based on their number. (b) Venn diagram depicting the distribution of top 50 ranked ligands across onco-fetal cell types they are associated to. (b) Heatmap of cell type-specific activity ($MUC6^{+}$ bipotent cells) for ligand-target pairs with high average scores in the neighborhoods harboring onco-fetal cells and $MUC6^{+}$ bipotent cells.

#### Supplementary Tables

Supplementary Table 1: Association of genes in deferentially active ligand-target pairs and corresponding communication domains (inferred for mouse brain dataset) with known anatomical brain regions (with supporting literature).

| Gene | Associated Communication Domain | Associated Region | Supporting Literature |
|---|---|---|---|
| *Ccnd1* | 13 | Amygdala | www.proteinatlas.org/ENSG00000110092-CCND1/brain |
| *Nr2f2* | 13 | Amygdala | www.proteinatlas.org/ENSG00000185551-NR2F2/brain |
| *Cort* | 0 | Cortex | www.proteinatlas.org/ENSG00000241563-CORT/brain |
| *Rspo1* | 9 | Cortex | www.proteinatlas.org/ENSG00000169218-RSPO1/brain |
| *Vip* | 5 | Cortex | elifesciences.org/articles/55130 |
| *Crlf1* | 16 | Hippocampus | www.proteinatlas.org/ENSG00000006016-CRLF1/brain |
| *Il16* | 17 | Hippocampus | www.proteinatlas.org/ENSG00000172349-IL16/brain |
| *Ntf3* | 16 | Hippocampus | learnmem.cshlp.org/content/13/3/307 |
| *Dlk1* | 3 | Hypothalamus | journals.plos.org/-plosone/article?id=10.1371/journal.pone.0036134 |
| *Tac1* | 14 | Hypothalamus | academic.oup.com/endo/article/156/2/627/2422577 |
| *Tgfa* | 7 | Striatum | www.jneurosci.org/content/8/6/1901.long |
| *Avp* | 15 | Thalamus | onlinelibrary.wiley.com/doi/10.1002/cne.22635 |
| *Lef1* | 2 | Thalamus | link.springer.com/article/10.1007/s00429-012-0474-6 |
| *Ntng1* | 2,10 | Thalamus | molecularbrain.biomedcentral.com/articles/10.1186/1756-6606-7-19 |
| *Tcf7l2* | 2,10 | Thalamus | link.springer.com/article/10.1007/s00429-012-0474-6; neuraldevelopment.biomedcentral.com/articles/10.1186/s13064-018-0107-8 |
| *Cntn2* | 1 | White Matter | www.proteinatlas.org/ENSG00000184144-CNTN2/brain; www.frontiersin.org/articles/10.3389/fncel.2019.00454/full |
| *Csf1* | 1 | White Matter | www.frontiersin.org/articles/10.3389/fimmu.2019.02199/full |

Supplementary Table 2: Association of top-ranked ligands and corresponding communication domains (inferred for mouse brain dataset) with known anatomical brain regions (with supporting literature).

| **Gene** | **Associated Communication Domain** | **Associated Region** | **Supporting Literature** |
|---|---|---|---|
| *Bdnf* | 16 | Hippocampus | https://www.nature.com/articles/4001957 |
| *Igf1* | 5 | Cortex | www.frontiersin.org/journals/cellular-neuroscience/articles/10.3389/fncel.2017.00014/full |
| *Vegfa* | 3 | Thalamus | link.springer.com/article/10.1007/s00018-013-1280-x |
| *Nrg1* | 1 | Cortex | www.sciencedirect.com/science/article/pii/S221112471400610X |
| *Npy* | 0 | Amygdala | www.nature.com/articles/s41467-024-49766-0 |
| *Nrg2* | 5 | Cortex | https://www.nature.com/articles/mp201722 |
